# Genome-scale metabolic model of the diatom *Thalassiosira pseudonana* highlights the importance of nitrogen and sulfur metabolism in redox balance

**DOI:** 10.1101/2020.10.26.355578

**Authors:** Helena M. van Tol, E. Virginia Armbrust

**Author notes:** Corresponding authors (HV), (EA).

## Abstract

Diatoms are unicellular photosynthetic algae known to secrete organic matter that fuels secondary production in the ocean, though our knowledge of how their physiology impacts the character of dissolved organic matter remains limited. Like all photosynthetic organisms, their use of light for energy and reducing power creates the challenge of avoiding cellular damage. To better understand the interplay between redox balance and organic matter secretion, we reconstructed a genome-scale metabolic model of *Thalassiosira pseudonana* strain CCMP 1335, a model for diatom molecular biology and physiology, with a 60-year history of studies. The model simulates the metabolic activities of 1,432 genes via a network of 2,792 metabolites produced through 6,079 reactions distributed across six subcellular compartments. Growth was simulated under different steady-state light conditions (5-200 μmol photons m^−2^ s^−1^) and in a batch culture progressing from exponential growth to nitrate-limitation and nitrogen-starvation. We used the model to examine the dissipation of reductants generated through light-dependent processes and found that when available, nitrate assimilation is an important means of dissipating reductants in the plastid; under nitrate-limiting conditions, sulfate assimilation plays a similar role. The use of either nitrate or sulfate uptake to balance redox reactions leads to the secretion of distinct organic nitrogen and sulfur compounds. Such compounds can be accessed by bacteria in the surface ocean. The model of the diatom *Thalassiosira pseudonana* provides a mechanistic explanation for the production of ecologically and climatologically relevant compounds that may serve as the basis for intricate, cross-kingdom microbial networks. Diatom metabolism has an important influence on global biogeochemistry; metabolic models of marine microorganisms link genes to ecosystems and may be key to integrating molecular data with models of ocean biogeochemistry.

## Introduction

Diatoms are unicellular photosynthetic eukaryotes derived from a secondary endosymbiotic event when a heterotrophic eukaryote engulfed a red algal cell and acquired a plastid [1]. They appeared in the fossil record ∼180 million years ago [2] and are distinguished from other photosynthetic organisms by their distinct combination of metabolic pathways, including the presence of a complete urea cycle, and their ability to precipitate silica to form their cell wall and to synthesize chitin. It is a special challenge of systems biology to understand how this unique combination of interlocking pathways has allowed diatoms to thrive in the dynamic conditions of oceanic ecosystems.

A central need for photosynthetic organisms is to balance redox reactions, particularly when nutrient availability limits growth (Figure 1). Light energy drives linear electron flow from water split at photosystem II (PSII) to PSI, generating reducing power (NADPH) and a proton gradient across the thylakoid membrane that drives ATP synthase. An ATP/NADPH ratio of 1.5 is required for CO_2_ reduction by the Calvin-Benson-Bassham cycle [3]. Linear electron flow alone has an ATP/NADPH ratio of ∼1.28 [4]. If not somehow mitigated, the resulting imbalance would cause the plastid to become over-reduced, damage the thylakoid membranes and cause photoinhibition [5]. In plants, NADPH-consuming pathways and alternative electron pathways that produce ATP without generating NADPH help balance redox reactions in the chloroplast [6,7]. Alternative electron fluxes include cyclic electron flow (CEF) around PSI, and water-to-water cycles where electrons from water oxidation at PSII are re-routed to an oxidation pathway – the Mehler reaction, chlororespiration, or photorespiration [8]. Diatoms are instead thought to preferentially regulate the ATP/NADPH ratio via energetic coupling between plastids and mitochondria [9], by which reduced metabolites are shuttled from the plastid to fuel ATP generation in the mitochondria.

**Figure 1.**
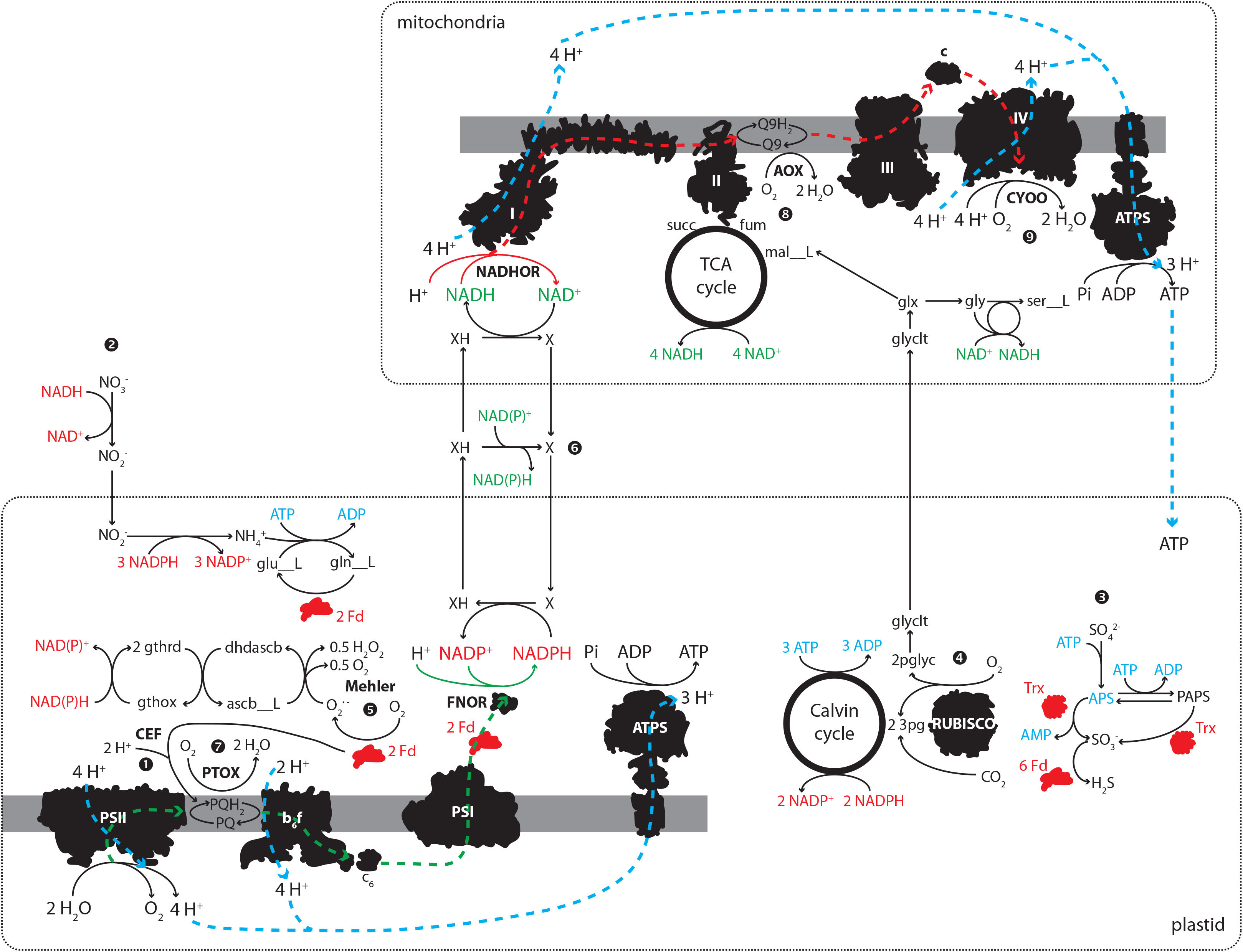
Diagram illustrating the principle reactions involved in generating ATP and balancing the ATP/NADPH ratio in diatom plastids. Black shapes: enzymatic complexes, gray bars: plastidial or mitochondrial membranes, dashed lines: grouping of plastidial and mitochondrial reactions, green: reactions that produce NAD(P)H equivalents, red: reactions that consume NAD(P)H equivalents, blue: ATP producing and consuming reactions. Abbreviations: L-glutamate (glu L), L-glutamine (gln L), 2-phosphoglycolate (2pglyc), glycolate (glyclt), glyoxylate (glx), glycine (gly), L-serine (ser__L), L-malate (mal L), fumarate (fum), succinate (succ), dehydroascborbate (dhascb), L-ascorbate (ascb L), oxidized glutathione disulfide (gthox), reduced glutathione (gthrd), reduced ferredoxin (Fd), reduced thioredoxin (Trx), plastoquinone (PQ), plastoquinol (PQH_2_), NADH ubiquinone oxidoreductase (NADHOR), ATP synthase (ATPS). Number symbols indicate reaction references in the text including: cyclic electron flow (CEF, ❶), nitrate assimilation ❷), sulfate assimilation (❸), ribulose-1,5-bisphosphate oxygenase (RUBISO, ❹), the Mehler reaction (❺), energetic coupling between the plastid and the mitochondria (❻), plastid terminal oxidase (PTOX, ❼), alternative oxidase (AOX, ❽), cytochrome *c* oxidase (CYOO, ❾).

Nitrate and sulfate assimilation are also involved in dissipating reducing equivalents in plastids as many enzymes involved in these processes are plastid-targeted (Figure 1). Nitrite reductase consumes 3 NADPH to reduce nitrite to ammonia, and the GS-GOGAT cycle (glutamine synthase – glutamine oxoglutarate aminotransferase) utilizes 2 reduced ferredoxin and 1 ATP to assimilate ammonia. During sulfate assimilation, sulfate is converted to APS (adenosine-5’-phosphosulfate) and either APS reductase or PAPS (3’-phosphoadenosine-5’-phosphosulfate) reductase utilizes a reduced thioredoxin to produce sulfite, consuming the equivalent of 2 ATP and 1 NADPH. Sulfite reductase consumes 6 reduced ferredoxin to produce sulfide for cysteine. A sulfurtransferase consumes 1 NADPH to produce L-cysteate from PAPS. Reductants are not consumed by the synthesis of sulfolipids from sulfite. Phosphate is assimilated by ATP synthase in the plastid and in the mitochondria. There is no evidence that diatoms can reduce phosphate to phosphite or phosphonate [10].

Metabolite production, secretion, or storage can also help balance redox reactions or dissipate energy [11], particularly when biomass production is otherwise inhibited. Phytoplankton adjust their biomass composition in response to nutrient limitation [12,13], elevated CO_2_ [14], low irradiance [15], and interactions with bacteria [16]. Diatoms also secrete more dissolved organic carbon in conditions of high light intensity [17], more dissolved organic nitrogen at suboptimal temperatures [18], and more exopolysaccharides (EPS) during nutrient limitation [19,20].

Here we created a mechanistic model of metabolism for the diatom *Thalassiosira pseudonana* CCMP 1335 to evaluate the interplay between redox balance, altered biomass composition, and organic matter secretion. We distilled all available physiological and molecular data from the literature to construct the metabolic network, to create biomass objective functions, to calculate ATP maintenance costs, and to add appropriate constraints to major fluxes. The model *i*Tps1432 includes the first mechanistic model of electron transfer by fucoxanthin chlorophyll *a/c* binding proteins (FCPs); it is also the first metabolic model to include silicate frustule formation and a hypothetical pathway for the biosynthesis of 2,3-dihydroxypropane-1-sulfonate (DHPS), a novel diatom osmolyte [21]. To simulate growth under a range of different light and nutrient conditions, we used Flux Balance Analysis (FBA) to calculate fluxes through the metabolic network given a set of constraints and an objective function to optimize [22]. We used biomass composition and photosynthetic production rate measurements from chemostats maintained under three different light levels [23] to construct light-dependent biomass objective function, constrain growth, photosynthesis, and respiration to calculate ATP maintenance costs and simulate metabolic fluxes under a range of irradiances. We found that cyclic electron flow could be an important sink for electrons at the light levels tested (5-200 μmol photons m^−2^ s^−1^), even though it is known to be a relatively minor component of alternative electron flow at higher light levels [9]. Nitrate reduction is also an important electron sink under these conditions. Next, we simulated growth in a nitrate-limited batch culture with dynamic Flux Balance Analysis (dFBA) using experimental biomass composition and PSII flux measurements from Liefer, *et al.* [24,25]. We found that sulfate reduction takes over the role of nitrate under these conditions. When biomass production is inhibited due to nutrient limitation, redox imbalances

## Materials and methods

### Network reconstruction and curation

A genome-scale metabolic model of *Thalassiosira pseudonana* CCMP 1335 was generated using *i*LB1027_lipid (the model of *Phaeodactylum tricornutum* CCAP 1055/1, [26]) as a starting point, based on similarities between the two diatoms. The *T. pseudonana* nuclear proteome was acquired from a dataset produced by Gruber, *et al*. [27], in which previous open reading frames (ORFs) were improved by ensuring that each gene starts with ‘ATG’, encodes an uninterrupted reading frame that ends in a stop codon, is less than 10 kb in length, and has EST support. The ORFs used in a *T. pseudonana* network reconstruction for BioCyc (including plastid and mitochondrial proteomes) were retrieved as well [28]; these proteins were re-annotated in 2012 using the JGI annotation pipeline for eukaryotes [29]. The *P. tricornutum* chromosomal proteome was downloaded from EnsemblProtists (ASM15095v2), the plastid and mitochondrial proteomes were downloaded from NCBI (acc no.: NC_008588.1, HQ840789.1). The authors of *i*LB1027_lipid provided a gene ID conversion table that we used to update the gene IDs in the published model. OrthoMCL [30] was used to identify gene orthologs of *P. tricornutum* and the two sets of *T. pseudonana* ORFs. A network of *T. pseudonana* reactions was generated by retaining reactions in *i*LB1027_lipid that contained gene orthologs from *T. pseudonana* and deleting reactions with no gene ortholog, except for spontaneous reactions. Protein localization predictions were performed on both *T. pseudonana* ORF sets (see Subcellular Protein Localization, below). BiGG IDs [31] were used for reactions and metabolites in the *T. pseudonana* network, and these were assigned to one of six different compartments: cytosol (‘c’), mitochondria (‘m’), peroxisome (‘x’), plastid (‘h’), thylakoid lumen (‘u’), or endoplasmic reticulum (‘r’).

We implemented the guidelines from an established genome-scale reconstruction protocol [32] to refine the *T. pseudonana* model. All genes from Gruber, *et al*. [27] with orthologs in the reconstruction, all genes assigned to a reaction in BioCyc [28], and all *T. pseudonana* genes without an ortholog in *P. tricornutum* were annotated with InterProScan [33] and those annotations were used to verify gene-protein-reaction associations, and to detect missing genes, reactions, and pathways in the model. We used KEGG [34] and BioCyc [28] databases to aid in model curation and to make comparisons between organisms. The literature on *T. pseudonana* was examined for experimental evidence for the existence of different reactions and for protein localization data (see references and notes in S1 Data Set). When experimental protein localization data was available, it superseded the subcellular localization prediction. For each reaction, mass and charge balance were verified and links to external databases were added for each reaction and metabolite. TransportDB 2.0 [35] was used to generate a list of transporter gene annotations; transport reactions were added to the model in cases where the database included associated substrates with the annotation. Extracellular transport reactions were also added in cases where there is experimental evidence that a substrate is excreted or utilized by *T. pseudonana* or other diatoms (see references and notes in S1 Data Set).

Dead-end metabolites are metabolites present only in blocked reactions; blocked reactions cannot carry flux due to reactions missing in the network and dead-end metabolites cannot be produced by the model. These reactions were identified with the flux analysis module in COBRApy and the gapfilling module was used to identify gaps in the network [36]. Gaps were filled if a gene for the missing reaction could be identified, if there is physiological evidence that the reaction exists, if the majority of the pathway was otherwise present in the model, or if the reaction was required to produce biomass. In *T. pseudonana*, we first checked whether the reaction was present in another compartment and if there was any evidence that the subcellular localization prediction was too stringent (e.g., a low confidence prediction may be more likely based on the localization of other reactions in the pathway), if there is the possibility of dual-targeting, or if a different gene with the correct localization for the reaction could be identified. Occasionally, the JGI ORFs (rather than the ORFs from the Gruber proteome [27]) provided the missing gene for a reaction, or a more likely subcellular localization prediction. If there was no evidence that a protein in the model was incorrectly targeted, then transport reactions between compartments were added to connect the network. To identify and remove erroneous energy-generating cycles (EGC), we followed the method proposed by Fritzemeier, *et al.* [37], by using the GlobalFit algorithm [38] to suggest the minimum number of changes required to remove an EGC. The algorithm found that the reaction transporting water between the cytosol and the mitochondria (‘H2Ot_m: h2o_c <=> h2o_m’) and the ITP-apyrase reaction (‘ITPA_c: h_c + idp_c + pi_c --> h2o_c + itp_c’) created energy-generating cycles. EGCs were removed by setting the lower bound of H2Ot_m to zero and by setting the upper bound of ITPA_c to zero.

Broddrick, *et al.* [39] published a list of twenty-nine modifications to the *P. tricornutum* model in 2019. We evaluated whether these modifications should also apply to *i*Tps1432 and adapted our model accordingly (S3 Data Set). *i*Tps1432 is available in SBML format as S1-S3 Files and has been deposited in the BioModels database (acc no.: MODEL2010230001-3).

### Subcellular protein localization

The protein localization pipeline developed by Levering, *et al.* [26] was updated for the *T. pseudonana* analysis. For plastid targeting predictions, TargetP [40] was replaced with ASAFind [27], a plastid proteome prediction tool developed for diatoms and other algae with plastids derived from a secondary endosymbiosis. All *T. pseudonana* proteins were used as input for SignalP 4.1 [41], TargetP 1.1 [40], HECTAR 1.3 [42], Mitoprot II 1.101 [43], ASAFind 1.1.7 [27], predictNLS 1.3 [44], and scanProsite [45]. PredictNLS, a tool for predicting nucleus targeted proteins, was run in batch mode using a script that re-implements predictNLS 1.3 in Python (https://github.com/peterjc/pico_galaxy/tree/master/tools/predictnls). ScanProsite was run to search for two peroxisomal targeting signals “[SAC]-[KRH]-[LM]>” and “S-S-L>” [46] and the PROSITE pattern PS00342 describing microbody C-terminal targeting signals, as well as the endoplasmic reticulum (ER) targeting signal “[KD]-[DE]-E-L>” [40,47] and the PROSITE pattern PS00014 describing other endoplasmic reticulum targeting sequences. All other programs were run with default settings. ER-targeted proteins were defined as ‘Not plastid, SignalP positive’ or ‘Plastid, low confidence’ by ASAFind and contained an ER-targeting signal identified by scanProsite. Plastid-targeted proteins include those identified by ASAFind as Plastid, high confidence’ or ‘Plastid, low confidence’ with no recognized ER targeting signal. Mitochondria targeted proteins are SignalP negative and predictNLS negative and match one of the following criteria: (A) have a Mitoprot II score > 0.9, (B) have a Mitoprot II score > 0.8 and are mitochondria targeted according to HECTAR or have a mitochondrial targeting peptide according to TargetP, or (C) are predicted to be mitochondria targeted by HECTAR and have a mitochondrial targeting peptide according to TargetP. Peroxisome targeted proteins are SignalP negative, predictNLS negative, not mitochondria targeted, and contain a peroxisome targeting signal according to scanProsite. Proteins in the plastid and mitochondrial genomes were assigned to reactions in the plastid and mitochondria, respectively. All remaining proteins were assigned to the cytosol. A total of 408 sequences from the optimized gene catalog could not be run through this subcellular localization pipeline despite curation by Gruber, *et al.* [27] because they have internal stop codons or were either too long or too short for some of the programs.

### Mechanistic model of light-harvesting

Using the *Synechococcus elongatus* model *i*JB785 [48] as a guideline, stoichiometric reactions were generated to represent light harvesting in *T. pseudonana*. The pigment weight and composition [49,50] of cells acclimated to each light-level were used in combination with the weight-specific absorption spectra for each pigment [51,52] to calculate the relative absorption of each pigment within 20 nm bins in the photosynthetically active radiation range (PAR range; 400-700 nm). Excitation energy transfer reactions were generated to account for energy loss in the transfer of excitation energy from different pigments in the fucoxanthin-chlorophyll *a/c* binding proteins (FCPs) to chlorophyll *a* in the reaction centers. Photon absorption was constrained for each wavelength using the absorption spectrum of *T. pseudonana* cells acclimated to different light levels [53–55] and the light intensity spectrum of a cool white fluorescent bulb according to the methodology in Broddrick, *et al.* [48]. We also extended the PSI and PSII reactions to include charge separation and recombination, as in *i*JB785. Photodamage of the D1 subunit was included as a component of the PSII reaction [48,56], and the metabolic cost of D1 repair was included as part of a non-growth associated ATP maintenance reaction as ATP-cost of phosphorylation and activation of the FtsH protease [57] and as ATP- and GTP-costs of biosynthesizing a D1 peptide. See S3 Data Set for calculations and references.

### Flux Balance Analysis

Flux balance analysis (FBA) simulates the flow of metabolites through a network of reactions using the mass balance equation,

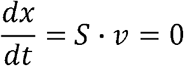

where the change in metabolite concentration (*dx*) over time (*dt*) is equal to the stoichiometric matrix (*S*) describing a reaction network times a vector of fluxes (*v*), given that

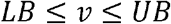

Intracellular metabolites are assumed to be at steady state, which allows *v* to be solved while maximizing an objective function within lower and upper bounds (*LB*, *UB*).

We used parsimonious Flux Balance Analysis (pFBA) to generate single solutions for each simulation. pFBA optimizes the model objective and then minimizes the total sum of flux to obtain a single solution. The second objective imitates a possible cellular objective of minimizing protein biosynthesis. The default GLPK solver in COBRApy [36] was used to optimize all FBA problems.

### Simulation of steady-state light limitation

pFBA was used with the solver tolerance set to 1e-8 to simulate growth of *i*Tps1432 in three chemostats maintained at three different light levels (5, 60, 200 μmol photons m^−2^ s^−1^) using photosynthetic production measurements from Fisher & Halsey [23] as constraints. The delivery rate of each nutrient was calculated based on the nutrient concentration in the media reservoir (0.25 mM NO_3_, 0.050 mM PO_4_, 0.106 mM Si(OH)_4_, 28.8 mM SO_4_), the chemostat cell concentration, the gram dry weight per cell calculated from biomass composition, and the chemostat dilution rate. Cultures were continuously bubbled to avoid carbon limitation; we assumed the ambient CO_2_ levels to be 390 ppm pCO_2_ and calculated the resulting concentrations of CO_2_ and HCO_3_ in the media [56]. Michaelis-Menten parameters were calculated from the iterature on *T. pseudonana* for CO_2_ and HCO_3_ [58]. The PSII reaction was constrained by fitting the Platt equation [59] (where *P_s_* and α are parameters of the hyperbolic tangent) to each gross photosynthesis (*GPP*) curve generated by Fisher and Halsey ([23], data provided by K. Halsey) and calculating the 95% confidence interval of gross photosynthesis at each light level.

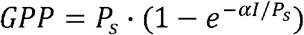

Photon absorption flux (PFA) was constrained for each 20 nm in the PAR spectral range using the method published in Broddrick, *et al.* [48],

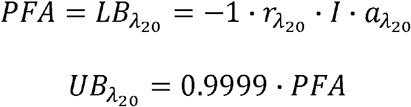

where the term 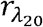 is the fraction of the light source’s photon flux at a particular wavelength integrated within a 20 nm bin, *I* is the light intensity (μmol photons m^−2^ s^−1^), and 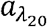 is the absorption of light at that wavelength (m^2^ gDW^−1^) [53–55]. The linear relationship between irradiance and D1 protein damage in *T. pseudonana* [56] was used to calculate the number of inactivation events per photon at each light level. The quantum efficiency values measured by Fisher & Halsey [23] were used to convert the number of inactivation events per photon to events per oxygen molecule evolved at PSII (S3 Data Set).

For these simulations, sink reactions for chrysolaminarin and glyceraldehyde-3-phosphate were added to the model to simulate the respiration of a transient carbon pool. Utilization of these pools was calculated as the difference between gross and net carbon production [23]. Measurements of dissolved organic carbon were used to constrain a symbolic expression representing the sum of organic carbon secretion. Light dependent respiration and mitochondrial maintenance respiration were calculated by Fisher & Halsey for each chemostat [23]. Light dependent respiration is defined as the difference between ^18^O_2_ signals in the light and the dark and includes all light-driven respiration reactions (the reduction of oxygen to water). We constrained the sum of all respiration reactions with a symbolic expression in which the lower bound is equal to the measured light dependent respiration values. Mitochondrial maintenance respiration is defined as the difference between gross carbon production and net oxygen production (converted to C units) and includes all organic carbon driven respiration reactions. We constrained the sum of all CO_2_ producing dehydrogenase reactions in the mitochondrial TCA cycle with a symbolic expression in which the lower and upper bounds are equal to the 95% confidence intervals of calculated mitochondrial maintenance respiration values. Jupyter notebook of this analysis can be viewed at https://github.com/hmvantol/diatom-FBA-notebook.

### Simulation of N-starvation

A dynamic FBA simulation was set up with the solver tolerance set to 1e-8 to simulate the progression in a batch culture of *T. pseudonana* from mid-exponential under NO_3_-limitation to mid-stationary phase under N-starvation using biomass composition and PSII photochemistry measurements from Liefer *et al.* [24,25]. Prior to the start of the experiment, cells were maintained at 85 μmol photons m^−2^ s^−1^ over a 12h: 12h light: dark cycle at 18°C in filtered seawater amended with half the concentration of all f/2 nutrients except for NO_3_, which was reduced by ~15 fold to 60 μM. At mid-exponential phase, the cells were diluted into fresh media with no added NO_3_ [24,25] and measurements were taken 0, 1, 3, 7, and 10 days after the dilution to “N-free” media, corresponding to mid-exponential, late-exponential 1, late-exponential 2, early stationary, and mid-stationary growth phases, as defined in Liefer *et al.* [24]. Nutrient concentrations in the seawater used for the media were not presented. We assumed a background seawater nutrient concentration of 20 μM NO_3_, 0 μM Si(OH)_4_, and 1.86 μM PO_4_. This assumption fit well with most nutrient measurements made 1 day after transfer to the new media and with estimates of nutrient concentrations in surface seawater at Cape Tormentine (Canada) from the World Ocean Atlas (~28 μM NO_3_, ~47 μM Si(OH)_4_, ~2 μM PO_4_).

To simulate the ten-day experiment, the growth period was divided into eighty 3-hour intervals. Steady-state was assumed for each time interval and pFBA was used to solve for the growth rate and reaction flux distributions every 3 hours. Units were converted from mmol to μmol prior to the pFBA step to avoid numerical precision issues associated with small fluxes. Each time point of the simulation involved four major steps: (1) calculate the biomass objective function, (2) calculate the uptake rates for each metabolite or nutrient and set photosynthetic constraints, (3) solve the pFBA problem, (4) update biomass composition and the environment.

The biomass objective function (*bof*) used at for each time interval was determined by calculating the relative difference between the simulated biomass composition at the current time point (*t_1_*) and the next biomass composition measurement (*t_m_*) provided by J. Liefer.

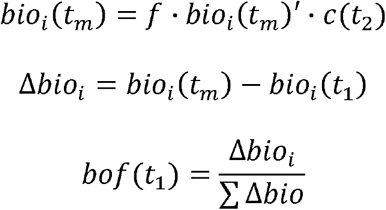

where each measured biomass component *i* is translated from pg/cell (*bio_i_* (*t_m_*)’) into g/L (*bio_i_* (*t_m_*) using cell abundance measurements (*c(t_2_)*), and typically *f* = 1. During the dark period, pigment biosynthesis was not included in the biomass objective function since *T. pseudonana* lacks a light-independent protochlorophyllide oxidoreductase [60]. Chrysolaminarin and triacylglycerides were also excluded from the dark period biomass objective function because these biomass components are known to be consumed at night [61]. Production of these compounds was increased during the light period by setting *f* to the fold-change values measured by Jallet, *et al.* in *P. tricornutum* grown across a 12: 12-h light: dark cycle [61]. During time points where the all ∆b;,i components are < 0, the objective function was switched to the ATP maintenance reaction (ATPM). At each time point, the upper bound of ATPM was re-calculated for changes in mg chl *a*/gDW using the *r_NGAM_* equation [62]. Individual pigment components (total chlorophyll, chlorophyll *a*, and total carotenoid) were also measured over the course of the experiment [25]. Data from the literature shows that the metabolites making up bulk biomass components shift in relative abundance over the course of N-starvation and during growth in batch culture. These were similarly tracked and used to as targets to construct new biosynthesis reactions for each time point. Protein [63] and free amino acid [64] composition measurements from nitrogen-replete and nitrogen-starvation conditions were used to calculate biomass targets for each growth phase; nitrogen-replete composition was applied as a target for mid-exponential and late exponential phase 1, while nitrogen-starved composition was applied as a target to late exponential phase 2, early and mid-stationary phases. Membrane lipid [65] and triacylglyceride [66] composition was measured during exponential, transition, and stationary phases in nitrogen-limited conditions. We used these measurements to calculate biomass targets for each growth phase: exponential composition was used as a target for mid-exponential and late exponential phase 1; transition phase composition was used as a target for late exponential phase 2; and stationary phase composition was used as a target for early and mid-stationary phase. See S4 Data Set for all calculations and references.

Using the method described by Chiu *et al.* [67] and published by Noecker *et al*. [68], we determined the uptake limit for each metabolite *j* at each time point *t*, such that the lower bound of each exchange reaction is equal to either the flux predicted by the Michaelis-Menten equation or the concentration of each metabolite *x_j_* per gram dry weight (*gDW*) biomass (*bio(t)*) per time period ∆*t*, whichever is closer to zero.

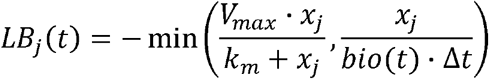

Nutrient uptake rates were calculated using Michaelis-Menten parameters from the literature on *T. pseudonana* for CO_2_, HCO_3_ [58], NO_3_ [69], NH_4_ [70], and PO_4_ [71] (S3 Data Set). The *V_max_* was set to 0.2 and *K_m_* was set to 0.05 for all other uptake reactions, with the exception of H_2_O and H^+^ which were assumed to diffuse freely and *V_max_* was set to 1000. The biomass-dependent lower bound assumes that *i*Tps1432 exists in a well-mixed environment where it can sense nutrient-availability and adjust its uptake rate accordingly [67].

PSII flux was calculated from fast repetition rate fluorometry (FRRf) parameters and culture data (electron transfer rate from PSII, PSII reaction centers per chlorophyll *a*) measured on days 0, 1, 3, 7, and 10 of the experiment (S3 Data Set, data provided by J. Liefer, [24]). The bounds of the PSII reaction were set as the 95% confidence interval of error propagated from the different measurements. To constrain the relative rate of CO_2_ assimilation to O_2_ production at each time point, we calculated the range of potential photosynthetic quotients (*Q*) by balancing the equation for the oxidation of new biomass to NO_3_ or NH_4_ with the chempy package [72]. Oxygen utilization and inorganic carbon evolution was not constrained during the dark period. After each time point, the concentration of O_2_, HCO_3_, and CO_2_ was re-equilibrated with the atmosphere so that concentrations remained constant (well-mixed). We included a constraint for photorespiration where the oxygenase flux of ribulose-1,5-bisphosphate carboxylase/oxygenase (RuBisCO) ranges from 0.001 – 0.025 the carboxylase flux.

For each time point, constraints were converted from mmol mg chl *a*^−1^ h^−1^ to mmol gDW^−1^ h^−1^ using simulated chlorophyll *a* and biomass concentration (mg chl *a* L^−1^/gDW L^−1^) ratios. Photon absorption flux was constrained for each time point as before, assuming a light absorption spectrum for cells acclimated to medium light [54] and the light intensity spectrum of a cool white fluorescent bulb (S3 Data Set). A new set of photon absorption (PHOA) reactions were constructed for each 20 nm wavelength at each time point using the simulated pigment composition.

We allowed for the mobilization of the biomass components chlorophyll *a*, chitin, polyphosphate, chrysolaminarin, protein amino acids, free amino acids, RNA, and triacyglycerides by including sink reactions for each of these components. The availability of each component was calculated as follows if ∆*bio_j_* is < 0.

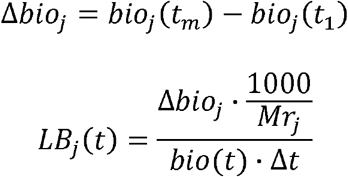

The degradation pathway for chlorophyll *a* is not characterized in diatoms, but chlorophyll *a* degradation was required to approximate the mg chl *a* gDW^−1^ factor. A demand reaction for chlorophyll *a* was included in the simulation and its lower bound was set to −1*∆*bio_j_*.

We adapted the method described by Chiu *et al.* [67] to calculate changes in biomass composition and the concentration of metabolites or nutrients over time. To calculate biomass composition, we calculated biomass concentration *bio(t_2_)* for the next time interval ∆*t* using the exponential growth equation,

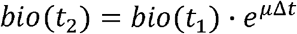

where μ is biomass demand or *v_DM_biomass_c_*. Next we calculated the fraction of new biomass attributed to each component using the biomass objective function. To calculate the concentration of each metabolite *x_j_* in the next time interval ∆*t*,

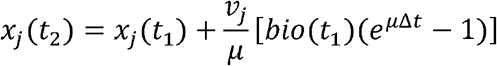

where *v_j_* is the flux of each metabolite, and *bio(t_1_)* is the biomass density or sum of all components at the current time point. Scripts for this simulation can be obtained from https://github.com/hmvantol/diatom-dFBA.

## Results

### Reconstruction of the *Thalassiosira pseudonana* metabolic model

A genome-scale metabolic model of *T. pseudonana* CCMP 1335 (*i*Tps1432) was generated using as a framework the *Phaeodactylum tricornutum* (CCAP 1055/1) models *i*LB1027_lipid [26] and *i*LB1034 [39]. Our model is distinct from a smaller, recently generated model of *T. pseudonana* (*i*Thaps987) designed to explore production of industrially useful compounds [73]. Differences between *i*LB1027_lipid and *i*LB1034 and their relevance to *i*Tps1432 are listed in S3 Data Set. *i*LB1027_lipid has more reactions than *i*LB1034 due to the inclusion of detailed lipid metabolism. Additional elements incorporated into the *T. pseudonana i*Tps1432 model include changes in the subcellular localization of reactions based on protein targeting sequences, improvements in modeled light absorption, and the inclusion of known metabolic differences between the diatoms. Several blocked reactions were resolved in *i*Tps1432, and the number of dead-end metabolites in the network was reduced when compared to the *P. tricornutum* models. The *T. pseudonana* model contains 6,073 reactions that represent a network of 2,789 metabolites and 1,432 genes, approximately 10-12% of the *T. pseudonana* genome (Table S1).

The *i*Tps1432 model contains reactions localized to six compartments representing the cytosol, mitochondria, plastid, thylakoid lumen, peroxisome, and the extracellular environment. As with the *P. tricornutum* model, our prediction pipeline did not localize any metabolic reactions to the endoplasmic reticulum, although other proteins were localized there. We evaluated the accuracy of our protein localization prediction pipeline by comparing our predictions to proteomics data from mitochondria and plastid fractions isolated from *T. pseudonana* [74]. Because the resulting fractions were not pure, we used peptide counts to determine if a protein was enriched in the mitochondrial or plastid fraction or in the cell lysate. A protein was considered enriched if it made up a greater proportion of peptides in one fraction versus the others. Using this criterion, 200 proteins were enriched in the mitochondria, 190 proteins in the plastid, and 564 proteins in the cytosol or some other organelle. Overall, 46.5% of proteins enriched in the mitochondria and 57.4% of proteins enriched in the plastid were accurately localized by our prediction pipeline (Figure S1a, S1b). About 70% of the 563 proteins that were enriched outside of the plastid and mitochondria were assigned to other compartments by our pipeline (Figure S1c); one protein was predicted to be localized in the endoplasmic reticulum and seven were predicted to be localized in the peroxisome, suggesting that most proteins enriched in the cell lysate were probably from the cytosol. Genes encoding 258 of the proteins detected by Schober, *et al*. were present in the model prior to model curation based on the literature and a little over half of the proteins (146 out of 258) were accurately localized by the subcellular localization prediction pipeline. After curation, 269 of the proteins detected by Schober, *et al.* were present in the model (Figure S1).

Two major metabolic differences distinguish the two diatom species. First, *T. pseudonana* has an absolute requirement for vitamin B_12_ as it possesses only the B_12_-dependent methionine synthase gene METH [75]. Adenosylcobalamin, methylcobalamin, and aquacobalamin were included in the biomass reaction as estimated millimole proportions of 1-gram dry weight ([76], S2 Data Set), and the cofactors were included in the B_12_-dependent reactions. Second, *T. pseudonana* produces chitin [77]; *i*Tps1432 was modified to include chitin biosynthesis and degradation pathways.

An orthoMCL comparison [30] of homologous genes between *T. pseudonana* and *P. tricornutum* guided additional modifications to *i*Tps1432. These analyses confirmed that *T. pseudonana* lacks the enzymes guanine deaminase, tryptophanase, ATP citrate synthase, and β-carbonic anhydrase [78]. The citrate synthase reaction in *i*Tps1432 was modified to deprotonate water rather than phosphorylate ADP, and a cytoplasmic carbonic anhydrase was added to the model [79]. The analysis also detected a few other differences between *P. tricornutum* and *T. pseudonana*. Reactions present only in *P. tricornutum* were eliminated from *i*Tps1432; these include the isomerization of xylose to xylulose, transamination of 4-aminobutyrate, lysis of O-acetyl-L-homoserine to L-homocysteine and acetate, L-tryptophan deamination, agmatine hydrolysis, formamide hydrolysis, and guanine deamination. Reactions present only in *T. pseudonana* include chitin synthesis and hydrolysis, and cleavage of pyruvate into acetyl-CoA and formate.

We also extended the *i*Tps1432 model to include synthesis and respiration of the carbohydrate storage molecule chrysolaminarin, synthesis and hydrolysis of polyphosphate, exopolysaccharide biosynthesis, silicic acid condensation to a silica frustule, and pathways for the biosynthesis and respiration of 2,3-dihydroxypropane-1-sulfonate (DHPS) [21,80]. Lipid metabolism was re-configured in *i*Tps1432 to reflect known differences in lipid composition [65,66]. We included transport reactions for those amino acids known to be excreted at high concentrations [81]. If genes encoding putative transporters for amino acids were identified, then additional transport reactions for chemically similar amino acids were added, with the assumption that these amino acids could use the same transporter. We included transporters for the following amino acids: L-glutamate, L-aspartate, L-isoleucine, L-leucine, L-valine, L-asparagine, L-glutamine, L-alanine, L-histidine, L-serine, L-threonine, glycine, and L-proline. Additional transport reactions for biotin, D-lactate, cyanocobalamin, aquacobalamin, DMSP, DHPS, glycine betaine, N-acetyltaurine, formamide, formate, uracil, acetate, choline, xanthine, ATP, AMP, triphosphate, and UDP-N-acetyl-alpha-D-glucosamine, were added based on information from the literature, gene annotations, and information from other algal models [48,82].

An important addition to *i*Tps1432 was inclusion of a mechanistic model of photon absorption and electron transfer by fucoxanthin chlorophyll *a*/*c* binding proteins (FCPs). We included reactions that describe energy transfer efficiency from excited pigments to chlorophyll *a* in the photosystems, and pigment de-excitation reactions to dissipate excess energy as heat or fluorescence (see S3 Data Set). PSI and PSII reactions were modified to include charge separation and recombination reactions. Photodamage of the D1 subunit was included as a component of the PSII reaction [48,56], and the metabolic cost of D1 repair was included as part of the non-growth ATP maintenance reaction to calculate the ATP-cost of phosphorylation and activation of the FtsH protease [57] and the ATP- and GTP-costs of biosynthesizing a D1 peptide (S3 Data Set).

### Development of biomass objective functions

A critical step in creating genome-scale metabolic models is development of accurate biomass production reactions in which precursor metabolites are converted into the cellular components that comprise the millimolar contribution to 1 g dry weight (gDW) of cell mass under specific growth conditions [32]. In *i*Tps1432, the biomass reactions generate the DNA, RNA, protein, free amino acids, pigments, carbohydrates, lipids (phospholipids, sulfolipids, galactolipids, glycerolipids), triacylglycerides, chitin, chrysolaminarin, osmolytes, a silica frustule, polyphosphate, and a soluble pool of vitamins and cofactors that together define the total biomass of *T. pseudonana* under a particular environmental condition. Biomass reactions were developed for *T. pseudonana* cells acclimated to three different light levels (5, 60, 200 μmol photons m^−2^ s^−1^). Bulk biomass composition (carbohydrates, protein, total dry weight) and chlorophyll *a* concentration was measured in cells grown in chemostats maintained at 18°C under 5, 60 and 200 μmol photons m^−2^ s^−1^ [23]. Pigment composition was derived from cells grown in a photobioreactor maintained at 18°C at 30 μmol photons m^−2^ s^−1^ [49] and from exponentially growing cells maintained at 18°C at 83 and 237 μmol photons m^−2^ s^−1^ [50]. The remaining biomass components were calculated from the literature, typically from exponentially growing cells, at temperatures ranging from 15-21°C (optimal growth temperature is 21°C); DNA nucleotide composition was calculated from the genome sequence data. A simplified siliceous frustule formation reaction was added to the model. The number of condensation reactions per Si atom was derived from NMR data on the degree of silica hydroxylation/condensation in *T. pseudonana* [83] and was used to calculate how much water should be released per gram of frustule formed. The weight of the frustule was calculated based on the expected degree of hydroxylation in the frustule and on either a linear relationship between growth rate and Si/C under light-limiting conditions (5, 60 μmol photons m^−2^ s^−1^) or a power law relationship under N-limiting conditions (200 μmol photons m^−2^ s^−1^) [84]. Calculations and references are provided in S2 Data Set.

The biomass composition also impacts the modeled photon absorption rate. Photon absorption integrated over the Photosynthetically Active Radiation (PAR, 400-700 nm) spectrum in 20 nm units was calculated from whole cell absorption spectra for cells acclimated to 25, 40-60, and 250 μmol photons m^−2^ s^−1^ [53–55], as well as weight-specific absorption spectra [51,52] and pigment composition (S3 Data Set).

Development of these biomass objective functions yielded information about the composition of *T. pseudonana* under different conditions and thus potential growth and acclimation strategies. The resulting total cell dry weights (scaled to the radius of each circle in Figure 2) is inversely related to light intensity (22.4, 16.6, 17.8 pg/cell for 5, 60, 200 μmol photons m^−2^ s^−1^, respectively) Similarly, the silica frustule was the greatest component of biomass at the lowest light intensity (5.2 pg/cell vs 1.6 and 0.7 pg/cell, Figure 2), likely a consequence of the slower, light-limited divisions rates combined with non-saturable silicic acid uptake kinetics in *T. pseudonana* [85]. Protein contribution to biomass was inversely correlated with light intensity, with the least amount of protein at the highest light levels. In contrast, total carbohydrates increased with light intensity and represented the largest component at 200 μmol photons m^−2^ s^−1^ ([23], Figure 2). Pigments per cell were greatest at 5 μmol photons m^−2^ s^−1^, resulting in an increased rate of photon absorption compared to the higher light levels. Photoprotective pigments (β-carotene, diadinoxanthin, diatoxanthin) were most abundant at 200 μmol photons m^−2^ s^−1^ ([49,50], Figure 2).

**Figure 2.**
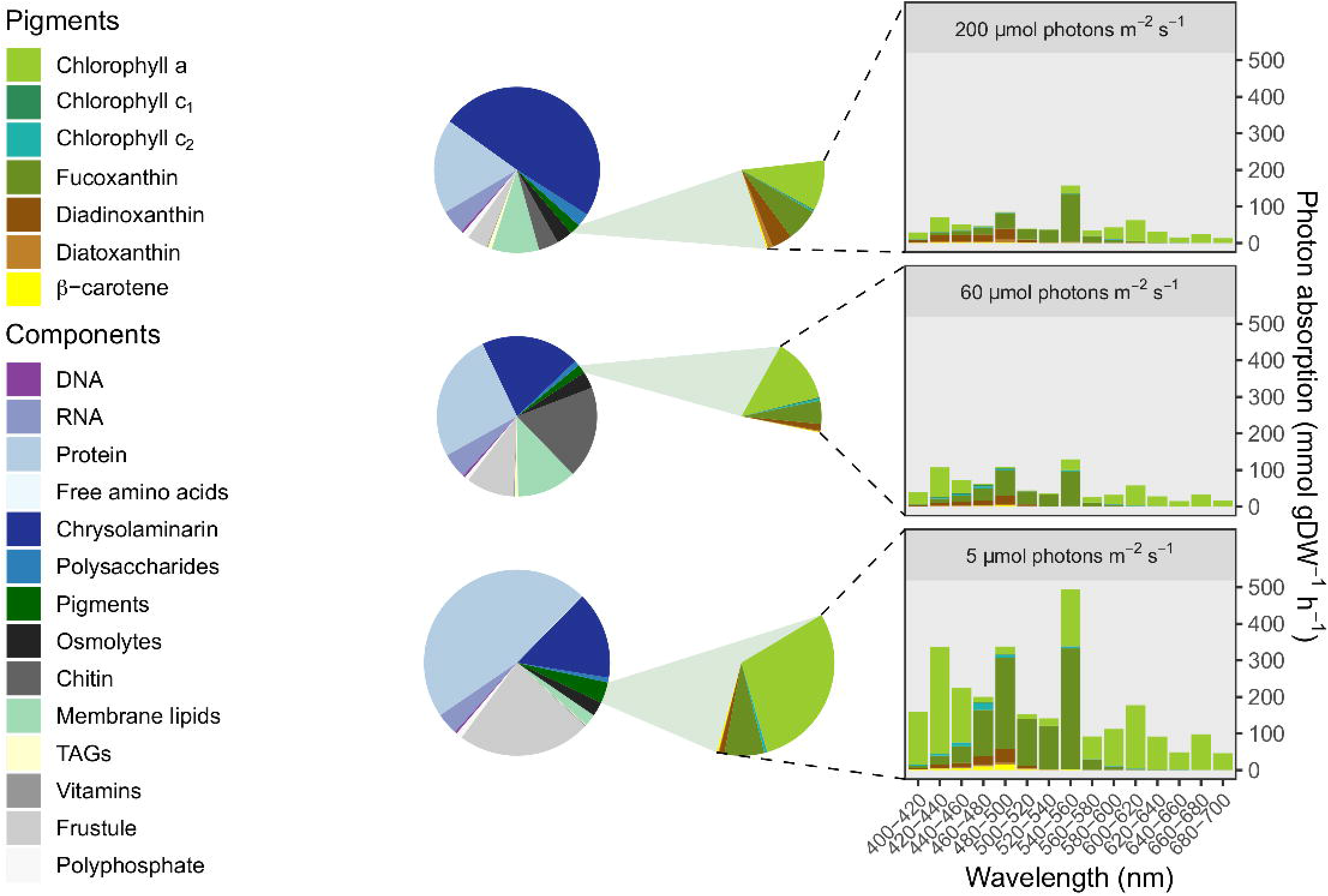
Biomass composition and detailed pigment composition of *Thalassiosira pseudonana* acclimated to three different light levels (5, 60, 200 μmol photons m^−2^ s^−1^). The radius of each circle is scaled to the total cellular dry weight. To demonstrate the effect of pigment composition (highlighted in green) on photon absorption at different wavelengths, photon absorption was plotted for each acclimated cell during illumination at 100 μmol photons m^−2^ s^−1^ under a cool white fluorescent bulb. The contribution of each pigment is integrated over each 20 nm of the light spectrum.

### ATP maintenance cost calculation

Cellular energy requirements, in the form of ATP utilization, impact biomass production and metabolite excretion in metabolic models. ATP maintenance costs were calculated for the three light levels (5, 60, 200 μmol photons m^−2^ s^−1^) using our biomass objective functions and production measurements (provided by K. Halsey) from chemostats maintained at the same light levels [23]. Growth-associated ATP utilization represents the energy not accounted for in biopolymer formation reactions while non-growth associated ATP utilization accounts for energy utilization in the absence of growth. Growth-associated ATP maintenance calculations can be impacted by photodamage due to high light intensities [86]. In the chemostat studies [23], the maximum photochemical yield of PSII (Fv/Fm) remained constant (0.56-0.58), across the three light levels indicating a lack of photodamage. We therefore constrained biomass production using chemostat dilution rates and performed Flux Balance Analysis (FBA) for each light level by maximizing the ATP hydrolysis reaction. A linear relationship (R^2^ = 0.94) between ATP utilization and growth rate allowed us to estimate the non-growth associated maintenance (*r_NGAM_* = −27 ± 52 SE) costs as the *y*-intercept and the growth associated maintenance (*r_GAM_* = 3809 ± 1221 SE) costs as the slope (S2 Figure). The small sample sizes resulted in large error bars (S2 Figure). Rather than constrain the ATP maintenance reaction with these bounds, we iteratively searched for individual *r_GAM_* values for each biomass objective function using the measured growth rate and calculated theoretical upper bounds for *r_NGAM_*.

The theoretical upper bound of non-growth associated maintenance (*r_NGAM_*) was calculated for each chemostat using the compensation light level (*I_c_*), which is the light intensity at which photosynthesis is equal to respiration and the growth rate is zero [62].

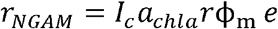

where *a_chla_* is the chlorophyll *a* specific absorption coefficient (m^2^ mg Chl *a*^−1^), *r* is the ratio of chlorophyll *a* to gram dry weight per cell (mg Chl *a*^−1^ gDW^−1^), *ϕ_m_* is the quantum efficiency of photosynthesis (mol O_2_ mol photon^−1^), and *e* is the amount of ATP generated per oxygen. The parameter I_k_ was estimated by fitting Chalker equation 1 [87] (where DR stands for dark respiration)

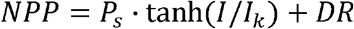

to each net photosynthesis (*NPP*) curve generated by Fisher and Halsey ([23], data provided by K. Halsey). *I_k_* was then used to calculate the compensation light level (*I_c_*) for each chemostat (Chalker equation 4).

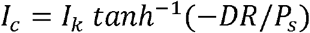

The *r* and *ϕ_m_* parameters were obtained from Fisher and Halsey; the *a_chla_* parameter was calculated from Finkel [53], Stramski *et al.* [54], and Sobrino *et al.* [55] (S3 Data Set). The resulting non-growth and growth associated maintenance costs are 1.6, 2.2, and 3.7 mmol gDW^−1^ h^−1^ and 2698, 2217, and 3669 mmol ATP gDW^−1^ for cells acclimated to 5, 60, and 200 μmol photons m^−2^ s^−1^, respectively.

### Cyclic electron flow

Cyclic electron flow (CEF) and respiration both consume the reduced ferredoxin and downstream equivalents generated via linear electron flow (LEF), driving ATP generation and increased flux through PSII (Figure 1) and thus balance redox. Additionally, CEF helps generates a proton gradient across the thylakoid membrane through cytochrome *b_6_/f*. Constraining these reactions can improve the accuracy of flux predictions in *i*Tps1432. Bailleul *et al.* [9] proposed that CEF is less than 5% of maximal total electron flow in diatoms. Maximal total electron flow occurs when light is saturating and increased flux of electrons cannot occur. To constrain CEF in *i*Tps1432 we simulated the Bailleul *et al.* [9] light response experiment with FBA using the biomass equation we developed and absorption spectrum acquired for cells acclimated to medium light intensity (60 μmol photons m^−2^ s^−1^; their cultures were acclimated to 70 μmol photons m^−2^ s^−1^). The maximal total electron flow through a cell acclimated to medium light was estimated by setting the level of light exposure to ~2000 μmol photons m^−2^ s^−1^ (saturating). PSII, oxygen exchange, and bicarbonate exchange reactions were constrained by the 95% confidence intervals of gross and net photosynthesis and carbon uptake, respectively, measured for cells acclimated to 60 μmol photons m^−2^ s^−1^ and exposed to 1952 μmol photons m^−2^s^−1^ [23]. The rate of D1 damage for the PSII reaction was calculated based on the number of inactivation events per O_2_ molecule evolved [56], using a quantum efficiency (*ϕ_m_*) of 0.056 mol O_2_ per photon [23] (S3 Data Set). Nutrient uptake rates were calculated using the Michaelis-Menten equation with f/2 nutrient concentrations. The upper bound of CEF was constrained with the following symbolic expression where CEF is set to 5% of total electron flow,

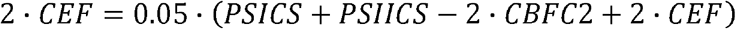

Linear electron flow (LEF) is the sum of electron flux through PSI and PSII (represented by charge separation reactions: PSICS, PSIICS) minus the linear flow from PSII through to PSI (represented by cytochrome *b_6_*/*f*: CBFC2). The pFBA simulation calculated 0.38 mmol gDW^−1^ h^−1^ as an upper bound of CEF and we used this value in subsequent simulations. CEF is relatively insensitive to short-term changes in light intensity and appears similar in a variety of diatom species [9]. At the highest light intensities (~2000 μmol photons m^−2^ s^−1^), CEF is an insignificant fractional component of total electron flow (TEF) in *T. pseudonana*; at the lower light intensity of ~100 μmol photon m^−2^ s^−1^, CEF corresponds to about half of TEF (see Extended Data Figure 9b in [9]). How CEF differs in cells acclimated to different light levels or cells in different growth phases is not yet known.

### Effect of light limitation

The allocation of photosynthetic energy to biomass production and respiration was quantified using pFBA with *i*Tps1432 at three different light levels [23], constrained by chemostat production data (provided by K. Halsey), the calculated ATP maintenance costs, and CEF, given the light-dependent differences in biomass composition (Figure 2). The pathways contributing to the dissipation of reducing equivalents generated by light energy include: CEF; respiration (ribulose-1,5-bisphosphate oxygenase, glycolate oxidase, plastid terminal oxidase, the Mehler reaction, alternative oxidase, cytochrome *c* oxidase); nitrogen assimilation (NO_3_ reductase, NO_2_ reductase, glutamate synthesis); sulfate assimilation (PAPS reductase, APS reductase, 2-aminoacrylate sulfotransferase, SO_3_ reductase); carbon assimilation and biosynthesis of reduced metabolites (Figure 1). Some pathways also dissipate the electrons generated by transient pools of organic carbon respired to CO_2_. For example, rapidly dividing cells preferentially use storage polysaccharides such as chrysolaminarin, while slowly growing populations dominated by cells in G1 phase preferentially use newly formed glyceraldehyde-3-phosphate (G3P) [88]. Additional sink reactions for chrysolaminarin and G3P were included to simulate respiration of these transient organic carbon pools. From the chemostat dilution rates we calculated the fraction of cells dividing per hour (*T. pseudonana* cell division is not synchronized). The sink reactions were constrained (proportionally to the fraction of cells dividing per hour) with the differences between gross and net carbon production (the carbon catabolism measurements) for each chemostat [23].

Biomass production and cyclic electron flow are the largest electron sinks (Figure 3a). G3P production by the Calvin cycle is a sink for electrons across all three light levels, with the lowest amount produced at the highest light intensity due to increased supply of G3P that is not entirely consumed by mitochondrial maintenance respiration. The biosynthesis of complex macromolecules, or further reduction of G3P, is another source of NADPH consumption, and becomes increasingly important at higher light intensities. Nitrate and sulfate reduction also consume reducing equivalents. At 5 μmol photons m^−2^ s^−1^, nitrate reduction is a proportionally more important component of biomass production than at other light levels because of the high protein content of cells acclimated to low light (10.5 pg/cell vs. 4.3 and 3.3 pg/cell, Figure 2). Respiration is also more important at low light levels. Mitochondrial maintenance respiration was used to constrain respiration of organic carbon via the TCA cycle, and as a result cytochrome *c* oxidase is a constant proportion of TEF (Figure 3a). PFBA predicts that the Mehler reaction was the largest respiratory flux at 5 and 200 μmol photons m^−2^ s^−1^, while at 60 μmol photons m^−2^ s^−1^ cytochrome *c* oxidase was the largest and there was also some respiration by plastid terminal oxidase (Figure 3a). To achieve a single solution, pFBA minimizes the absolute sum of fluxes with the objective of minimizing enzyme utilization. Given that this objective may not be relevant to photosynthetic organisms dealing with redox balance, we explored additional respiratory constraints in the simulation.

**Figure 3.**
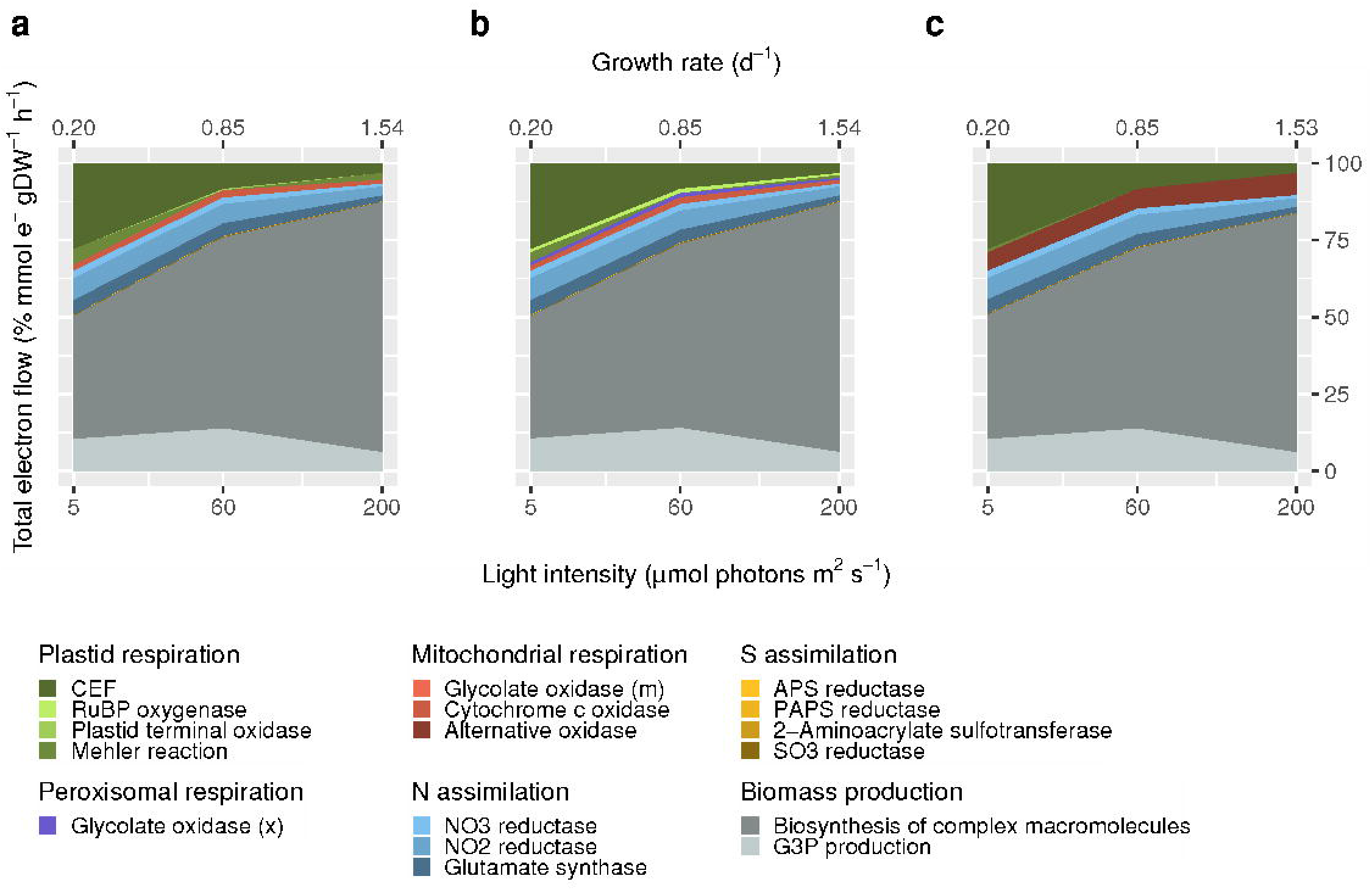
Parsimonious Flux Balance Analysis (pFBA) predictions (in mmol e^−^ gDW^−1^ h^−1^) for the contribution of different reactions to the dissipation of reductants generated by photosynthesis and organic carbon utilization in *i*Tps1432 across three chemostats maintained at 5, 60, and 200 μmol photons m^−2^ s^−1^. Differences in biomass composition and biosynthesis of metabolites impact the contribution of C, N, and S assimilation reactions (grey, blue, yellow, respectively). Respiration reactions are distributed across several different organelles including the plastid, peroxisome, and mitochondria (green, purple, red, respectively). Flux predictions with (a) baseline constraints on respiration, (b) photorespiration included, (c) photorespiration and energetic coupling included.

First, we noticed that photorespiration was not a part of the original pFBA prediction and that it could be an important sink for reductants in photosynthetic organisms. Photorespiratory flux, or the specificity of ribulose-1,5-bisphosphate carboxylase/oxygenase (RuBisCO) for CO_2_ versus O_2_, was calculated with the following expression [89]

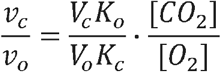

where *v_c_* is the carboxylase flux, and *v_o_* is the oxygenase flux through RuBisCO, *V_c_K_o_/V_o_K_c_* is the specificity factor of RuBisCO for CO_2_ over O_2_, and [CO2]/[O2] is the ratio of CO_2_ versus O_2_ in the pyrenoid. We used a specificity factor of 79, as determined for the related diatom *T. weissfloggii* [90] because of an assumed similarity in function based on the predicted peptide level similarities between the RuBisCO enzymes from the two diatoms (rbcS is 97% identical, rbcL is 98% identical). The concentration of CO_2_ in the pyrenoid, where most RuBisCO is located, is estimated at 100 μM [91]. The concentration of O_2_ in the pyrenoid is unknown and is difficult to measure [92]. We therefore used the ambient concentration of O_2_ in seawater at equilibrium with the atmosphere (200 μM).

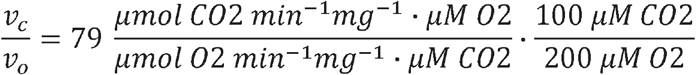

Photorespiration was constrained with the following symbolic expression,

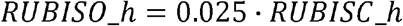

where the oxygenase activity of RuBisCO is 2.5% the carboxylase activity. The addition of a photorespiratory constraint activates the oxygenase activity of RuBisCO as well as peroxisomal glycolate oxidase (Figure 3b). For a K_m_ of 65 μM [90], pyrenoid CO_2_ concentrations are likely higher than estimated because diatoms are not carbon-limited, and O_2_ concentrations are thought to be lower (Young, pers. comm.) Thus, these simulations may overestimate the significance of photorespiration.

In diatoms, a major component of redox balance is the flow of reducing equivalents from the plastid to the mitochondria [9], based on the observation that flux of electrons through PSII depends on mitochondrial respiration. In a third experiment, we included energetic coupling between the plastid and the mitochondria by redirecting reductants generated by linear electron flow to NADH ubiquinone oxidoreductase with the following symbolic expression,

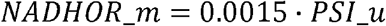

Little is known about the extent of energetic coupling in *T. pseudonana* and how it may change with acclimation to different light intensities, so the value of this constraint (NADHOR_m/PSI_u=0.0015) was chosen for exploratory purposes. Flow of reductants from the plastid activates the mitochondrial alternative oxidase reaction in *i*Tps1432 (Figure 3c). Larger values that increased the flux through alternative oxidase impacted the growth rate of *i*Tps1432 because there was a trade-off with CEF which appears to be required for energy generation at these lower light levels.

The simulations under different light limiting conditions highlighted the interconnected pathways that diatoms rely on to balance reductant dissipation through biomass requirements and different respiratory pathways. These results raise the question of how the dissipation of reductants generated by photosynthesis occurs under nutrient-limitation conditions when biomass production cannot be used as an electron sink.

### Effect of nitrogen starvation

The previous simulations highlighted the importance of nitrate reduction as an electron sink under steady-state conditions. We next explored the impact of nitrogen limitation and starvation on reductant flow using dynamic Flux Balance Analysis (dFBA) to simulate growth in batch culture where cells are not in steady-state. Throughout the simulation, we tracked simulated nutrient and excreted metabolite concentrations in the media as well as molecular and elemental biomass composition. The simulations relied on experimental data [24,25] derived from *T. pseudonana* cells grown at 85 μmol photons m^−2^ s^−1^ in a 12: 12h light: dark cycle at 18°C in f/2 media modified to initiate N-starvation. *T. pseudonana* acclimated to medium light levels (60 μmol photons m^−2^ s^−1^) is the closest condition examined in the previous simulation. Biomass composition and macro-nutrient concentrations were determined experimentally at 0, 1, 3, 7, and 10 days after the start of the experiment. Measurements of biomass C, N, and P were made more frequently at 0, 1, 2, 3, 5, 7, and 10 days after the start of the experiment. Liefer, *et al.* [25] defined four different growth phases according to observed growth patterns: mid-exponential corresponded to N-replete steady-state growth (day 0); late exponential corresponded to reduced growth after dilution to N-free media prior to stationary phase (day 1-3); early stationary corresponded to one day after cessation of cell division (day 7); and mid-stationary corresponded to five days after cessation of cell division (day 10).

In batch culture, biomass composition measurements are equal to the composition of cells from all previous time points plus newly synthesized biomass minus the re-mobilized biomass components. Biomass objective functions were computed for each time point by comparing the simulated biomass composition at the current time point to the experimentally measured biomass compositions at the different time points (target biomass composition) (Figure 4a). As nitrogen is depleted, *T. pseudonana* uses up protein and accumulates carbohydrates and lipids; there is also accumulation of DNA as cells stop actively dividing, a decrease in RNA and pigments per cell, and accumulation of residual P likely corresponding to polyphosphate storage ([24], Figure 4a). Experimental measurements and simulations of biomass composition produce C:N or N:P ratios that indicate cell composition was impacted by nitrogen starvation after 3 days in culture (Figure 4b). After 10 days in culture, measurements of C:P decreased due to polyphosphate increasing as a cellular component in stationary phase. There is a similar change in C:P in the simulated *i*Tps1432 biomass composition (Figure 4b). Small differences between measurements and simulated elemental ratios could be attributed to both error in the bulk biomass composition data taken from the literature to supplement the experimental measurements (osmolytes, chitin, vitamins), errors in the assumed compositional data making up the larger components (molecular composition of RNA, protein, free amino acids, protein, carbohydrates, EPS, and lipids), or over-production and under-production of various components at different points in the simulation (S3 Figure). Nevertheless, many measured biomass components were accurately simulated and the predicted timing of nitrogen starvation was accurate, giving us confidence that our model reflects the principle effects of nitrogen limitation and starvation in batch culture.

**Figure 4.**
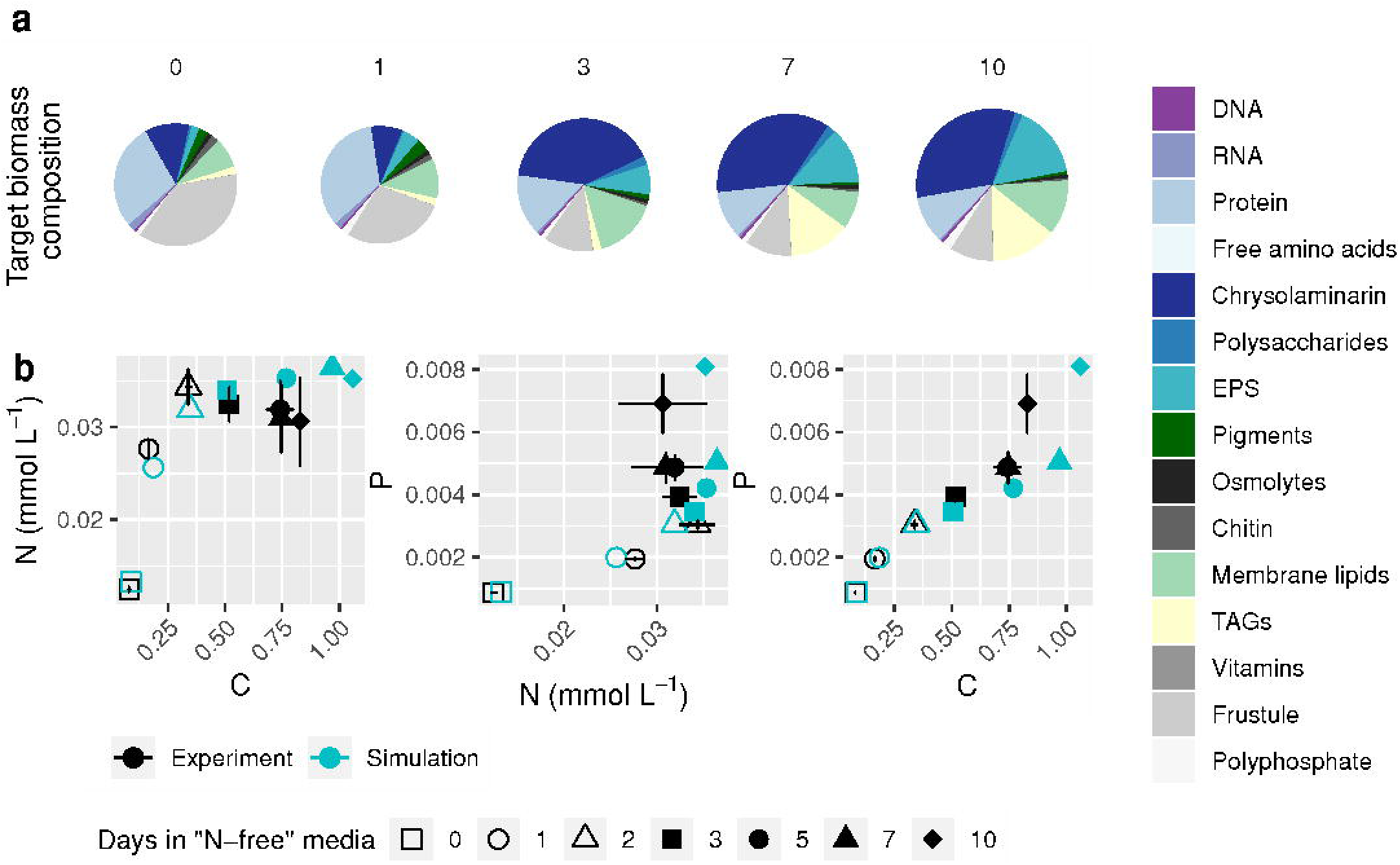
Target biomass composition and comparison between simulated and experimental biomass elemental ratios for the transition of *i*Tps1432 from exponential to stationary phase under nitrogen-limited conditions. (a) Biomass composition was measured 0, 1, 3, 7, and 10 days after transfer to fresh media with low nitrate and used to calculate the biomass objective function at each time point. (b) C:N, N:P, and C:P ratios were measured 0, 1, 2, 3, 5, 7, and 10 days after transfer and compared with simulated biomass elemental composition.

The added NO_3_ (and NO_2_) in the experimental media were depleted sometime between 1 and 3 days after the initiation of the experiment; low levels of background NH_4_ in the media were depleted after 1 day. Simulated NO_3_ was completely depleted from the media shortly after 1 day of growth whereas simulated NH_4_ was released into the media concurrent with the NO_3_ depletion and was subsequently taken up and re-excreted into the media between the different target days. The simulated PO_4_ concentration in the media matched the observed gradual decrease over time (Figure 5a). The greatest discrepancy between the measured and simulated nutrients was for Si(OH)_4_ concentrations. Experimental Si(OH)_4_ concentrations were simultaneously drawn down with the NO_3_ during the first day and then plateaued at ~10 μM. The simulated drawdown of Si(OH)_4_ instead plateaued at ~25 μM shortly after 1 day in culture. This difference likely reflects errors in the weight of the cell frustule used in the biomass objective function. This value was not experimentally measured and so the value used for the biomass objective functions was extrapolated from the difference between dry weight measurements and the sum of other biomass components (S4 Data Set). Inaccurate simulation of silicate utilization will have little impact on the flow of reductants as the silicate condensation reaction to form the frustule is not a redox reaction. To simulate a diel pattern of biomass formation, we included over-production of chrysolaminarin and TAGs during the light period in the biomass objective functions and did not include production of TAGs, chrysolaminarin, and pigments in the dark period [61]. Biomass production was strongest during the light period and was typically followed by respiration of some biomass components in the dark to create a diel pattern of changing biomass concentration (Figure 5b).

**Figure 5.**
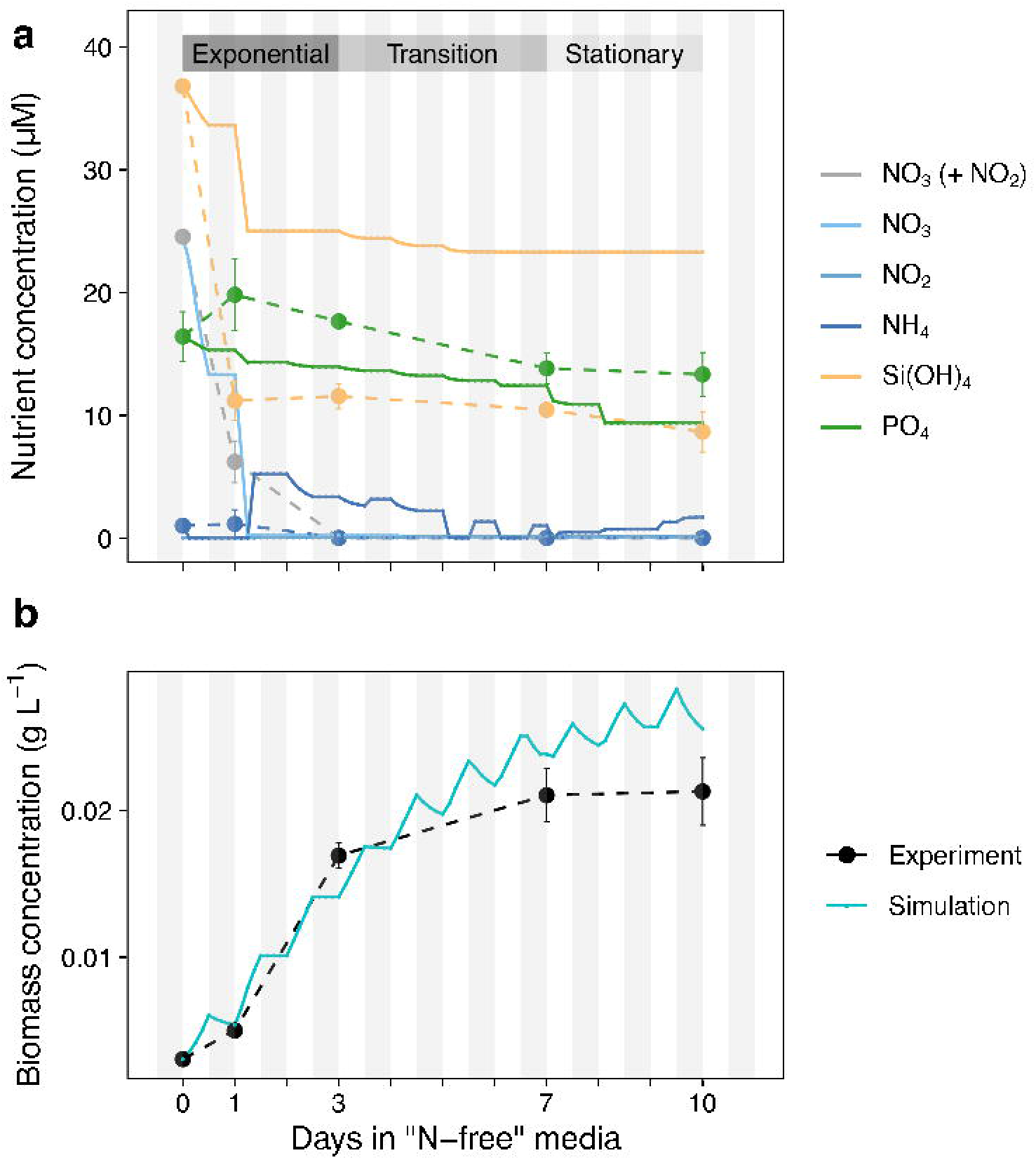
Comparison between simulated and experimental (solid, dashed lines) media nutrient concentration and biomass concentration for the transition of *i*Tps1432 from exponential to stationary phase under nitrogen-limited conditions. (a) NO_3_ (+ NO_2_), NH_4_, Si(OH)_4_, and PO_4_ were measured 0, 1, 3, 7, and 10 days after transfer to fresh media with low nitrate and compared to simulated media nutrient concentrations (b) Cell concentration (cells mL^−1^) was measured each day and converted to biomass concentration (g L^−1^) where measurements of cell dry weight were available and compared with the simulated biomass concentration. 12: 12 h light: dark cycles are depicted with white and grey stripes.

The ability to accurately simulate uptake of nitrate and C:N under non-steady-state conditions motivated a more detailed examination of electron flow. Gross oxygen evolution from the PSII reaction was constrained based on the electron transfer rate and the number of PSII reaction centers per chlorophyll *a* measured on days 0, 1, 3, 7, and 10 of the experiment [24]. Per milligram of chlorophyll *a*, gross oxygen evolution decreased slightly on the first day and remained steady until the seventh day, with a sharp increase on the tenth day of the experiment (Figure 6a). There is good agreement between gross oxygen production measured at 85 μmol photons m^−2^ s^−1^ for cells acclimated to medium light intensity grown in chemostats [23] to the PSII flux calculated with data from FRRf for cells in mid-exponential phase (Figure 6a). On a gram dry weight basis, there is an initial increase in PSII flux followed by a gradual decline over the course of the experiment as a result of declining chlorophyll *a* concentrations relative to total biomass. PSII flux is only slightly higher on the tenth day of the experiment on a gram dry weight basis (Figure 6a). The simulation includes diel fluctuations in chrysolaminarin and TAG production which create regular fluctuations in the ratio of O:C of new biomass (Figure 6b). The elemental formula of newly produced biomass was used to calculate a range of possible photosynthetic quotients (PQ: the proportion of oxygen evolved to inorganic carbon assimilated in the light) given growth on nitrate or ammonia (Figure 6c). The combination of constraints on the PSII, oxygen exchange, and inorganic carbon assimilation reactions limit respiration during the light period, while respiration is left unconstrained in the dark period. In general, total respiratory flux increases with increased flux through PSII, although there is some influence by the photosynthetic quotient (Figure 6c, 6d). Diel changes in respiration are controlled by diel patterns in biomass formation in which the rates of chrysolaminarin and TAG production vary throughout the light period. During the light period plastid terminal oxidase and the Mehler reaction are the primary respiratory fluxes. During the dark period, respiration switches to peroxisomal glycolate oxidase and cytochrome *c* oxidase as organic matter is respired for energy (Figure 6d).

**Figure 6.**
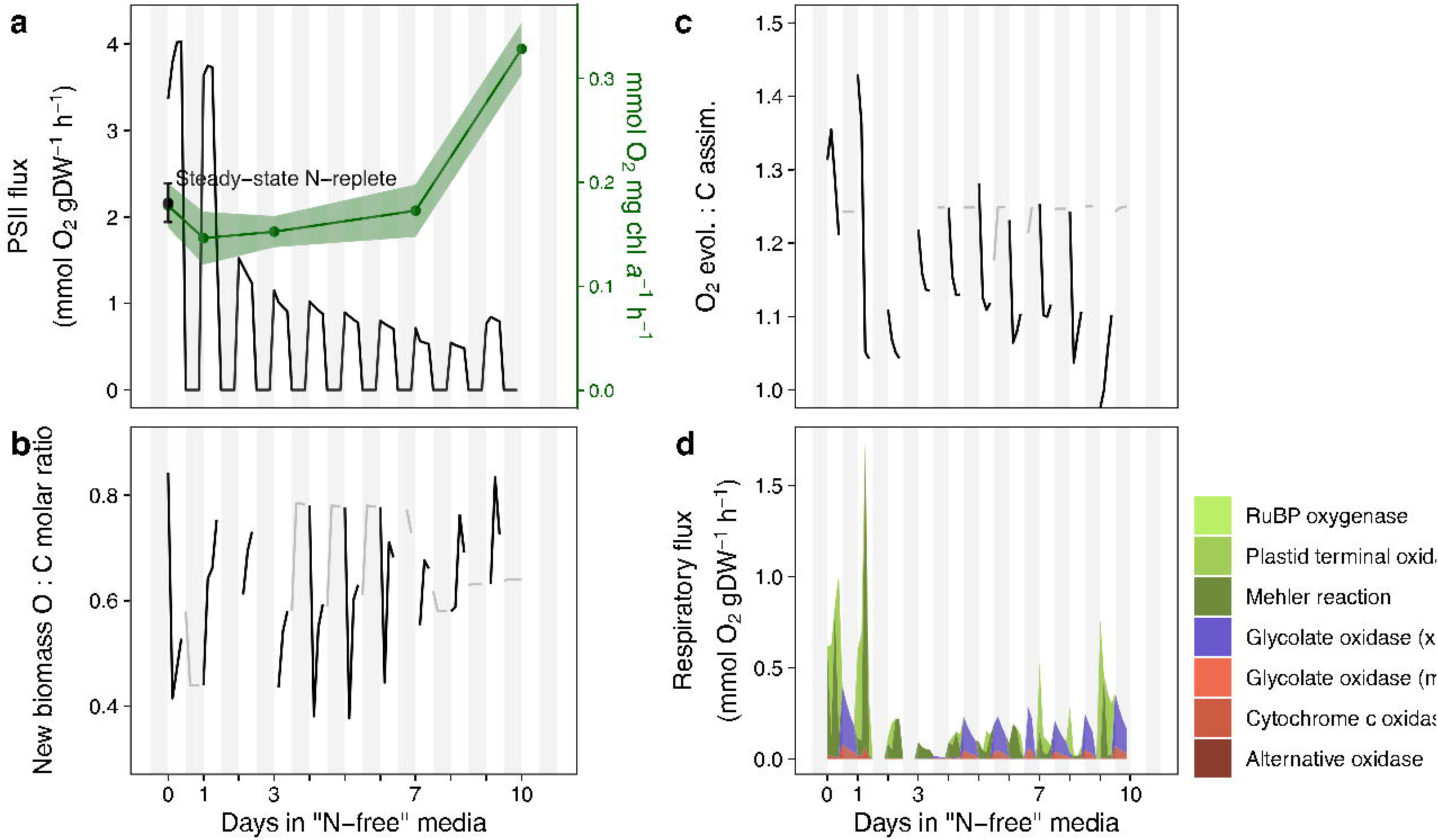
Simulated respiratory flux predictions (in mmol O_2_ gDW^−1^ h^−1^) and the constraints affecting respiration for the transition of *i*Tps1432 from exponential to stationary phase under nitrogen-limited conditions. (a) Flux through the PSII reaction is driven by photon absorption and is constrained by measurements taken 0, 1, 3, 7, and 10 days after transfer to fresh media with low nitrate (green, mmol O_2_ mg chl *a*^−1^ h^−1^, [24], S3 Data Set). The 95% confidence intervals were converted from mmol O_2_ mg chl *a*^−1^ h^−1^ to mmol O_2_ gDW^−1^ h^−1^ using simulated mg chl *a* / gDW and set as lower and upper bounds of the reaction. Steady-state N-replete ± 95% confidence interval (black) measurement of net oxygen production [23] was plotted at the 0 h time point. Simulation results are plotted here for each 3 h increment in mmol O_2_ gDW^−1^ h^−1^ (black). (b) Plot of simulated new biomass O: C molar ratio during the light (black) and dark (grey) period. (c) The photosynthetic quotient was calculated from the simulated new biomass elemental composition during the light period (black) and also contributes to the respiratory flux results. Oxygen and inorganic carbon assimilation were left unconstrained during the dark period (grey). (d) Respiratory flux predictions and contribution of different respiration reactions (plastid: green, mitochondria: red, peroxisome: purple). 12: 12 h light: dark cycles are depicted with white and grey stripes.

The photosynthetic quotient is the proportion of oxygen evolved to inorganic carbon assimilated in the light and its calculation assumes that all assimilated CO_2_ is used to generate biomass. However, diatoms are known to excrete dissolved organic carbon during photosynthesis. For this reason, we explored a range of different PQ constraints (95% PQ – 105% PQ) to see how this assumption impacts metabolite secretion (Figure 7). Metabolite excretion can occur if the by-product of a reaction cannot be assimilated by the organism; for example, cyanide is a by-product of cyanocobalamin utilization. Occasionally cyanide is used by 3-mercaptopyruvate sulfurtransferase in the de-sulfuration of 3-mercaptopyruvate, resulting in the excretion of thiocyanate. Cyanocobalamin is a form of vitamin B_12_ frequently used in culture media, but is not widely available in the environment. Polyphosphate is also a by-product in vitamin B_12_ metabolism and folate biosynthesis; excess is excreted and then re-assimilated at later time points as the preferred source of phosphorus (Diaz2019). In this simulation, metabolites are excreted predominantly as a result of biomass re-mobilization or to balance redox.

**Figure 7.**
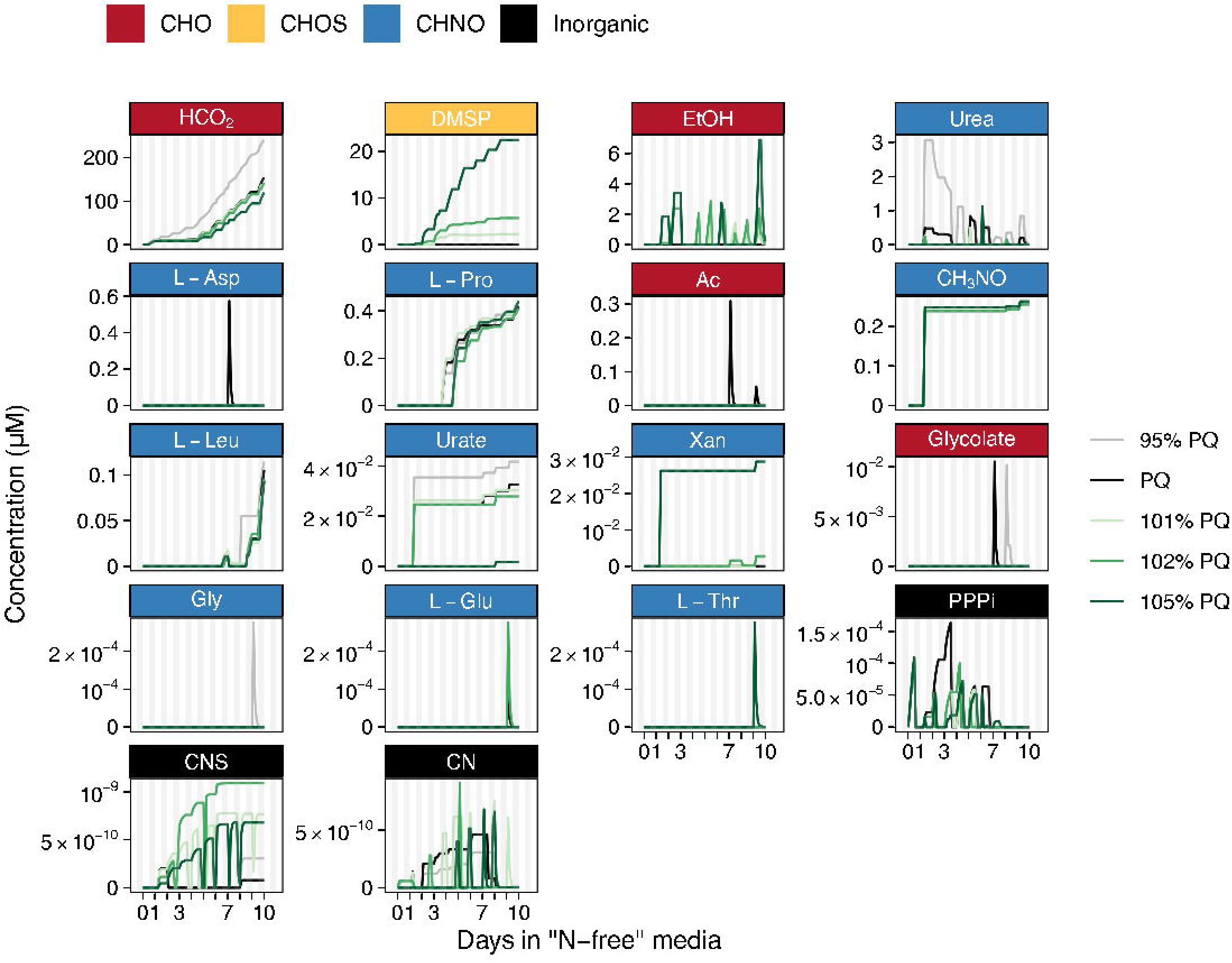
Simulated metabolite excretion by *i*Tps1432 during the transition from exponential to stationary phase under nitrogen-limited conditions across a range of possible photosynthetic quotient constraints. Note that *y*-axes are scaled to each metabolite and metabolites are listed in order of highest to lowest maximum concentration. Metabolites composed of organic carbon are labeled red, organic nitrogen are blue, organic sulfur are yellow, and inorganic are black.

Formate and DMSP production were most strongly impacted by changes in the PQ. DMSP production increases with the photosynthetic quotient (or when respiration is low). Conversely, formate production is highest when respiration is high (Figure 7). Similarly, more ethanol is excreted when the PQ is higher and more urea is excreted when the PQ is lower than predicted by biomass composition, and then both are re-assimilated at night. Other metabolites, proline, leucine, formamide, xanthine, and urate, appear to be by-products of re-mobilizing certain biomass components (protein, free amino acids, RNA) to achieve the targeted objective. Some metabolites including L-aspartate, acetate, glycolate, glycine, L-glutamate, and L-threonine appear to be more sporadically excreted and re-assimilated.

We evaluated reactions involved in N and S metabolism in simulations where DMSP is excreted and found increased sulfate assimilation after the onset of NO_3_ depletion (Figure 8). NO_3_ reductase uses 1 NADH to produce NO_2_ in the cytosol, NO_2_ reductase is localized in the plastid and uses 3 NADPH to produce NH_4_, and the GS-GOGAT cycle assimilates NH_4_ while utilizing ATP and 2 reduced ferredoxin when localized in the plastid. After NO_3_ is depleted, ammonia assimilation continues sporadically throughout the time course as a side-effect of the ammonia excreted due to the re-mobilization of different pools of organic nitrogen during biomass composition changes. SO_4_ can be assimilated in the plastid by a plastid-localized ATP:sulfate adenylyltransferase that produces adenylyl sulfate (APS). APS reduction to SO_3_ consumes a reduced thioredoxin, SO_3_ reductase is plastid localized and consumes 6 reduced ferredoxin to produce H_2_S. SO_4_ reduction to sulfide becomes more prevalent immediately after NO_3_ is depleted and results in DMSP excretion (Figures 7, 8). 2-aminoacrylate sulfotransferase also contributes to redox balance by consuming 1 NADPH in the production of L-cysteate which is thought to be an intermediate in DHPS production (Durham2019), a component of diatom biomass in *i*Tps1432.

**Figure 8.**
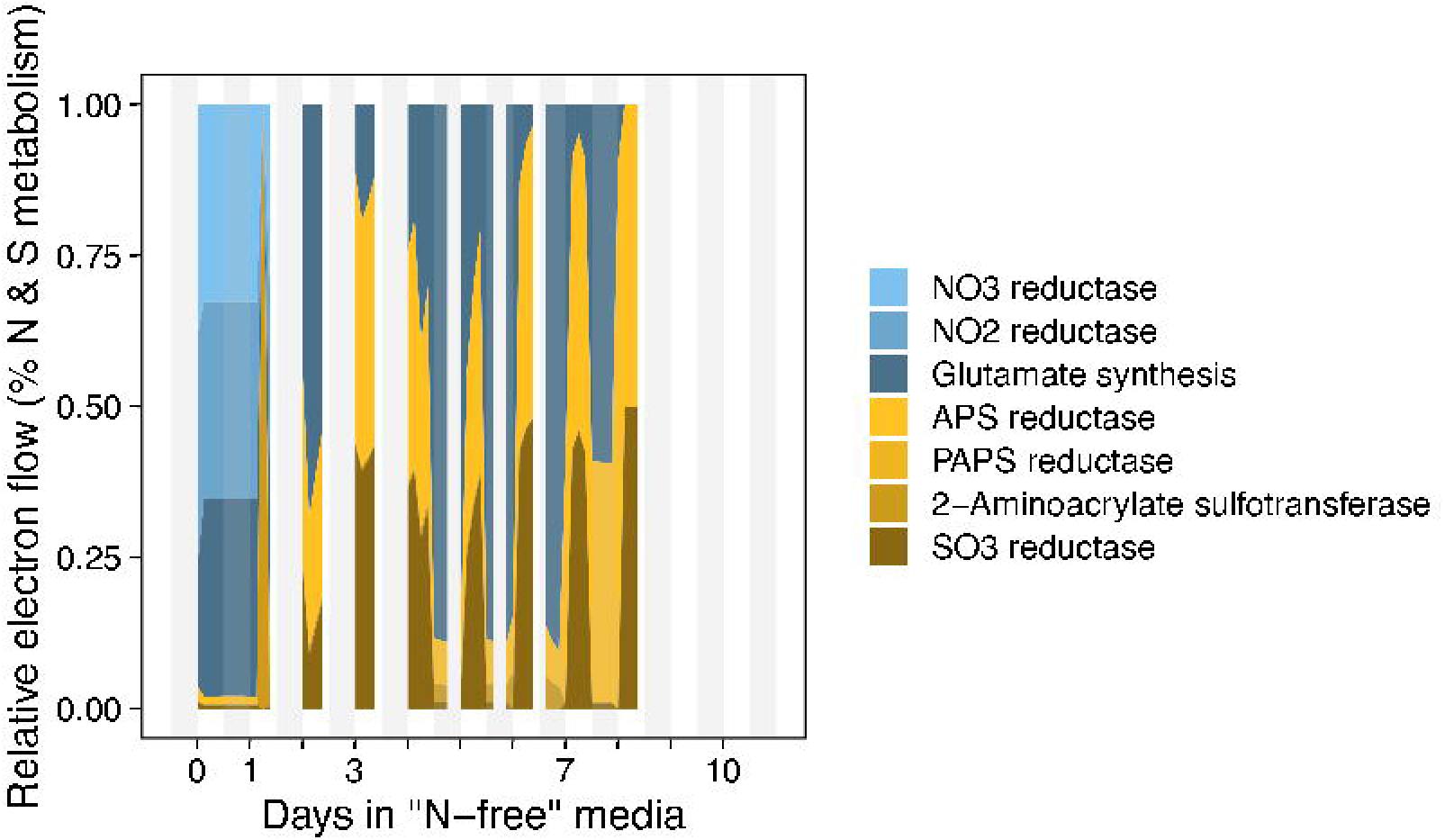
Simulated N & S (blue, yellow) metabolic flux predictions (in mmol e^−^ gDW^−1^ h^−1^, bottom panel) during the transition of *i*Tps1432 from exponential to stationary phase under nitrogen-limited conditions with a 102% photosynthetic quotient constraint.

## Discussion

The interdependence between biomass composition, photon absorption, nutrient utilization, and photosynthetic constraints is a distinguishing feature of *i*Tps1432, and a critical component of photosynthetic modeling given the dynamic nature of biomass composition and its role in photosynthesis. These interconnections allowed us to model the different ways that diatoms may maintain redox balance. A critical first step was development of biomass objective functions for three different light levels (5, 60, 200 μmol photons m^−2^ s^−1^) and use of chemostat production measurements to constrain flux for different light intensities (Figure 2). Given an elemental formula for biomass composition, the degree of reduction can be calculated as the number of electron equivalents per gram atom C [93]. The inverse relationship between protein content of cells and light meant that the degree of reduction was also inversely related to light (5.66, low light; 5.47, medium light; 4.97, high light). More highly reduced biomass composition as result of increased nitrate assimilation, in addition to increased light-dependent respiration at low light could be part of a previously proposed strategy that *T. pseudonana* limits respiration of organic carbon by using alternative redox balance strategies in order to improve the efficiency of biomass production [23].

Photosynthetic organisms have evolved a variety of mechanisms to deal with the primary challenge of photosynthesis: how to capture light energy while evading potential damage. Over longer time periods, diatoms adjust their pigment composition to deal with incoming light energy, but short-term fluctuations in light intensity with inadequate sinks for electrons could theoretically cause a plastid to become over-reduced and damaged. Energy dissipation as heat or fluorescence (non-photochemical quenching) is one potential mechanism, summarized in *i*Tps1432 as a photon loss reaction. The metabolic challenge of photochemical quenching is managing the mis-match between reduced ferredoxin produced by linear electron flow and the actual metabolic requirements for NADPH. Light-dependent respiration reactions in the plastid are one of the primary sinks for reductants. Alternatively, reduced metabolites can be exported from the plastid and oxidized by mitochondrial respiration. Finally, cyclic electron flow is another potential sink for reductants, as well as a non-reducing source of energy since the reaction drives proton pumping by cytochrome *b_6_/f*. Halsey & Fisher postulated that CEF could be more important at lower light levels, but did not measure it in their experiment [23]. Our FBA predicted that CEF is a major component of alternative electron flow, particularly at lower light levels (Figure 3a). This finding was unexpected as it has been shown that CEF is less than 5% of maximal electron flow and the transfer of reductants from the plastid to the mitochondria is more prevalent in diatoms when compared to plants [9]. The absolute flux of CEF remains relatively constant across all light levels and in multiple diatom species [9]. We used FBA to calculate an upper bound for CEF at saturating light levels (maximal electron flow). At low light levels, this flux value is relatively important given the decrease in total electron flow as light intensity goes down. In the data given by Bailleul *et al.* [9], CEF makes up about half of TEF at the lowest light level measured (~100 μmol photon m^−2^ s^−1^); so we may be underestimating its importance at these light levels due to the poorly constrained values of growth-associated ATP maintenance, and given the contribution of CEF to energy generation.

Respiration via cytochrome *c* oxidase is activated by the respiration of transient organic carbon pools by the mitochondrial TCA cycle. Some of the transient organic carbon that was calculated to be available (as the difference between gross and net carbon production measurements) is directly utilized, as evidenced by the decreased use of the Calvin cycle at 200 μmol photon m^−2^ s^−1^ (Figure 3a). Remaining respiratory flux was routed through the Mehler reaction or plastid terminal oxidase. In the Mehler reaction, oxygen reacts with reduced ferredoxin emerging from photosystem I and forms a superoxide anion. Superoxide dismutase neutralized two reactive anions into oxygen and hydrogen peroxide, and ascorbate peroxidase converts hydrogen peroxide into water. NAD(P)H is consumed through the glutathione-ascorbate cycle (Figure 1). Chlororespiration results from the oxidation of the plastoquinone pool by plastid terminal oxidase (Figure 1) and was historically defined as an electron transport chain in the thylakoid membrane involving a proton-translocating NADH:plastoquinone reductase complex (NDH1) and a plastid terminal oxidase (PTOX) [94]. *T. pseudonana* and other unicellular algae lack an NDH complex [95], and likely rely on a non-electrogenic NDH2 or ferredoxin:plastoquinone reductase (CEF) to fuel the terminal oxidase [94,96]. Chlororespiration is thought to be only active in low light conditions or darkness [96,97]. Here FBA predicts that PTOX is active at the medium light levels, and the Mehler reaction is active at low and high light. Parsimonious FBA minimizes the absolute sum of fluxes, and therefore returns a single solution with the lowest possible number of active fluxes. As a result, only one light-dependent respiration reaction is active and this may not be realistic. We experimented with different constraints on respiration using the available information in the literature (Figure 1).

Photorespiration occurs whenever there is oxygen in the pyrenoid because the enzyme RuBisCO cannot always distinguish between O_2_ and CO_2_. We activated photorespiratory flux by assuming a high concentration of O_2_ relative to CO_2_ in the pyrenoid. This constraint activated the oxygenase activity of RuBisCO as well as peroxisomal glycolate oxidase (Figure 3b). In diatoms, photorespiration is truncated and 2-phosphoglycolate is not recycled back to ribulose-1,5-bisphosphate [98]. Glycolate can be oxidized to glyoxylate and then converted to either malate or transaminated to glycine or is excreted under certain conditions [99,100]. Alternative oxidase was activated by imposing an energetic coupling constraint on PSI and NADH ubiquinone oxidoreductase to simulate energetic coupling [26] (Figure 3c). However, only a low level of energetic coupling (NADHOR/PSI = 0.15%) could be introduced before the constraint impacted biomass production. This observation supports the idea that CEF is more important at low light levels, and can be inhibited if reductants are diverted to the mitochondria. CEF helps generates a proton gradient in order to fuel ATP production; this could be an important reaction when light levels are low and cells are more energy starved, which is why our predictions are dependent on the GAM.

In addition to respiration, nitrate and sulfate assimilation also contribute to balancing redox reactions in the plastid, the relative impact of which depends on how cells adjust respiration rates in response to changing redox pressures [101]. We simulated the progression of a batch culture from nitrate limitation to N-starvation using dynamic FBA (Figure 5). When nitrate is depleted, diatoms continue to produce biomass by re-mobilizing internal sources of nitrogen and producing more of the carbohydrates [25]: chrysolaminarin and EPS (Figure 4a). Diatoms also experience decreased flux through PSII caused by pigment degradation [24], which decreases light absorption. Data on net oxygen evolution as nitrogen is depleted in batch culture was not available; we calculated a range of possible photosynthetic quotients from the new biomass equation at each time point based on biomass production from nitrate versus ammonia. This strategy only accounts for metabolites that are part of the objective function and doesn’t account for the possibility of excreted metabolites or the re-mobilization of biomass components. We tested a range of different PQ constraints (95% −105% the original PQ value) and found a potential role for sulfate assimilation after nitrate is depleted (Figure 8). Secretion of the organic sulfur compound DMSP as a result of increased sulfate assimilation is in line with previous experimental work indicating that nitrate limitation causes the greatest increase in intracellular DMSP production [102] (Figure 7). DMSP is hypothesized to act as an osmolyte replacing stores of proline, under nitrate limitation [102]. Many of the compounds produced and excreted by *i*Tps1432 are compatible solutes, a class of compounds known to be transported in and out of cells in response to changes in osmotic pressure. We did not account for the possibility of changing osmolyte composition as part of the biomass objective function as the only measurement of osmolytes are from N-replete conditions [21].

Experimental support for a role for nitrate uptake in response to fluctuating redox pressures comes from the observation that nitrogen-replete diatoms secrete ammonium during rapid increases in irradiance [103]. Additionally, enzymes within nitrogen and sulfur assimilation pathways in diatoms, are redox-sensitive [104]. Nitrate assimilation is likely to be more effective at dissipating reductants than sulfate assimilation because of low consumption of ATP relative to NADPH equivalents. However, sulfate is consistently available at high concentrations (28 mM, [105]) throughout the ocean whereas nitrate is typically limiting in both coastal regions [106] and in the subtropical gyres [107]. Although our focus was on growth limitation by nitrate, other types of nutrient-limitation (such as silicon, iron, zinc, or vitamin B_12_ [108]) may impact redox balance by limiting biomass formation.

In our simulations, *i*Tps1432 reduces ATP demand by limiting flux through the TCA cycle and instead secretes ethanol and formate (Figure 7), both of which are suggestive of some level of fermentation. Anaerobic bacterial cultures are known to accumulate fermentation products, and metabolic models of *E. coli* quantitatively predict their secretion [11]. Formate is a by-product of the methionine salvage pathway or DMSP biosynthesis, and is produced via pyruvate formate lyase, an alternative reaction to pyruvate dehydrogenase. In *E. coli*, ethanol is typically produced via alcohol dehydrogenase that detoxifies acetaldehyde generated by pyruvate decarboxylase during fermentation. *T. pseudonana* lacks the gene for pyruvate decarboxylase, and instead produced acetaldehyde as a by-product of threonine aldolase. In *i*Tps1432, acetaldehyde is either converted into ethanol and excreted or into acetate which may rejoin the TCA cycle as acetyl-CoA. Both cyanobacteria and the green alga *Chlamydomonas* accumulate fermentation products under dark anaerobic conditions [109,110], likely due to inhibition of mitochondrial respiration [111]. The diatom *P. tricornutum* appears to increase rates of mitochondrial respiration under stressful conditions (high light intensity, iron or nitrate limitation, or supraoptimal temperatures) as suggested by upregulation of alternative oxidase expression [112]. While *T. pseudonana* displays here the potential to secrete fermentation products, fermentation is likely minimal during normal culture conditions. We hypothesize that fermentation may instead occur in nature when respiration rates do not increase rapidly enough to match increased redox pressure such as may occur under fluctuating light conditions or in organisms with very rapid growth rates [113].

Amino acids are commonly secreted by a variety of diatoms [20,114]. We were surprised to predict the secretion of amino acids by *i*Tps1432 under nitrate starvation conditions (Figure 7). Secretion of these amino acids occurs mostly during shifts to new biomass targets where protein re-mobilization is possible and are not impacted by changes in respiration. We did not include transport reactions for small peptides although these compounds are an important component of secreted metabolites in *T. pseudonana*, possibly as a by-product of protein turnover [115]. During N-starvation we would expect protein re-mobilization to contribute to biomass formation for other metabolites requiring nitrogen rather than result in excretion of amino acids. This observation is similar to the degradation of RNA nucleotides into urate and xanthine and the excretion of formamide (Figure 7). Perhaps these nitrogenous compounds are too highly reduced to be useful during nitrate starvation and are therefore released into the environment.

Most of the metabolites secreted by *i*Tps1432 can be consumed by marine bacteria. For example, a subset of marine bacteria can utilize glycolate as a sole carbon source [116], and bacterial transcripts for glycolate oxidase were found to vary on a diel cycle during a phytoplankton bloom [117]. Many bacteria from the Roseobacter clade rely on organic nitrogen and sulfur compounds produced by phytoplankton or other bacteria as they are unable to reduce nitrate or nitrite and some cannot reduce sulfate [118]. Alphaproteobacteria and Gammaproteobacteria are known to degrade DMSP into methanethiol (CH_4_S) or dimethyl sulfide (DMS) [119]. Differences in the metabolic networks of phytoplankton as well as differences in respiration, and how different species react to fluctuations in environmental conditions and redox imbalances will impact the character and quantity of metabolites secreted. These are likely major factors that structure the bacterial community associated with phytoplankton and control bacterial succession over the course of a bloom [120].

## Conclusion

*Thalassiosira pseudonana* CCMP 1335 was isolated from Moriches Bay, New York, in 1958, and the whole genome was sequenced in 2004 [121]. With availability of the genome, *T. pseudonana* has been studied from a systems-wide perspective using transcriptomics, proteomics, and metabolomics (eg. [115,122–125]). The genome-scale metabolic model of *T. pseudonana* created here builds on previous modeling work [26], incorporates currently available physiological and genomic data, and will serve as a powerful tool to generate hypotheses about diatom metabolism and to interpret future experiments.

*i*Tps1432 could be extended in a variety of directions in the future. Reactions describing complex formation for metal- and cofactor-requiring proteins could be added to the model to better describe vitamin and trace metal utilization, as trace metal limitation is an important nutrient condition in the ocean [108] and vitamins play an import role in interactions with bacteria [126,127]. Additionally, an effort to better characterize transporters would significantly improve prediction of metabolite secretion and mechanisms of energetic coupling between the plastid and the mitochondria. The development of representative marine metabolic models, such as *i*Tps1432, could allow us to integrate molecular data with models of ocean biogeochemistry in the future.

## Supporting information

S1 Data Set

S2 Data Set

S3 Data Set

S4 Data Set

## Acknowledgements

Thanks to H.V.’s committee members, Elhanan Borenstein, Jody Deming, and Anitra Ingalls, for their insight and advice. Thanks to Kimberly Halsey & Justin Liefer for providing and discussing data from their publications. This work was supported by Gordon and Betty Moore Foundation grant GBMF3776 awarded to E. Virginia Armbrust. This document has been approved for release: LLNL-JRNL-815904.

## Supporting information captions

**Table S1** Comparison of attributes of *i*Tps1432 and *P. tricornutum* GEMs

**S1 Figure** Intersection between proteins enriched in the (a) mitochondria, (b) plastid, and (c) cell lysate (Schober, *et al.*, 2019), subcellular protein localization pipeline predictions, and proteins included in the model.

**S2 Figure** Relationship between measured growth rate and optimal ATP utilization in *i*Tps1432, given the biomass composition at different light levels. The *x* error bars represent the 95% confidence interval of measured dilution rates, and the *y* error bars follow from the 95% confidence intervals of bulk biomass measurements (carbohydrates, protein, total dry weight, in pg C/cell).

**S3 Figure** Comparison of simulated biomass components versus target components at the onset of different growth phases (days, 0, 1, 3, 7, 10). The best and worst (blue, red) estimates of biomass composition were marked for each biomass component. (A) Bulk biomass components, (B) RNA nucleotides, (C) Protein amino acids, (D) Free amino acids, (E) EPS sugars, (F) Pigment molecules, (G) sulfolipids, (H) phosphtidylcholines, (I) phosphatidylethanolamines, (J) phosphatidylglycerols, (K) diacylglycerides, (L) monogalactosyldiacylglycerides, (M) digalactosyldiacylglycerides, (N) triacylglycerides, (O) mg chl *a* /gDW.

**S1 Data Set** Table of reactions and metabolites and references in *i*Tps1432. (XLS)

**S2 Data Set** Biomass composition calculations and references for *Thalassiosira pseudonana* acclimated to 5, 60, and 200 μmol photons m^−2^ s^−1^. (XLSX)

**S3 Data Set** Constraint calculations and references for *Thalassiosira pseudonana*. (XLSX)

**S4 Data Set** Biomass composition calculations and references for *Thalassiosira pseudonana* during nitrogen-starvation

**S1 File** *i*Tps1432 acclimated to low light intensity (5 μmol photons m^−2^ s^−1^) in SBML format. Level 3, Version 1, fbc ver. 2. (XML)

**S2 File** *i*Tps1432 acclimated to medium light intensity (60 μmol photons m^−2^ s^−1^) in SBML format. Level 3, Version 1, fbc ver. 2. (XML)

**S3 File** *i*Tps1432 acclimated to high light intensity (200 μmol photons m^−2^ s^−1^) in SBML format. Level 3, Version 1, fbc ver. 2. (XML)

**Table S1.**
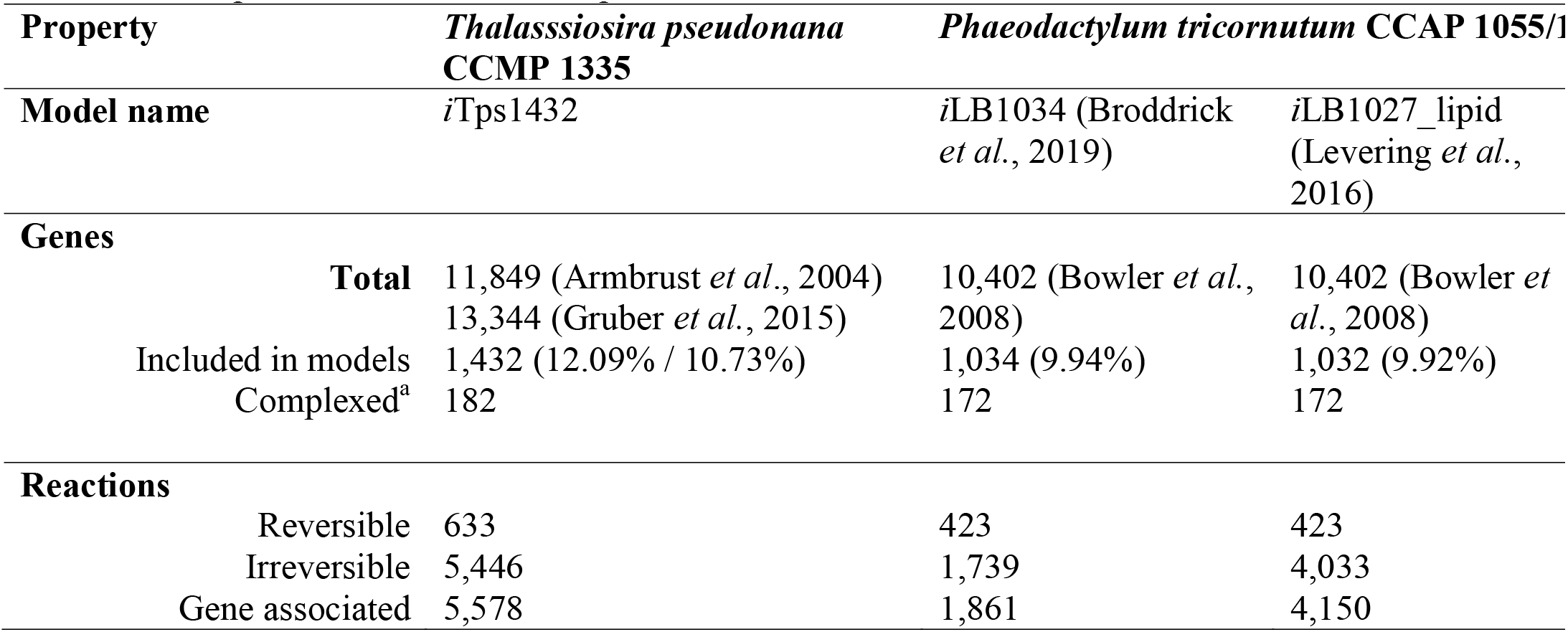
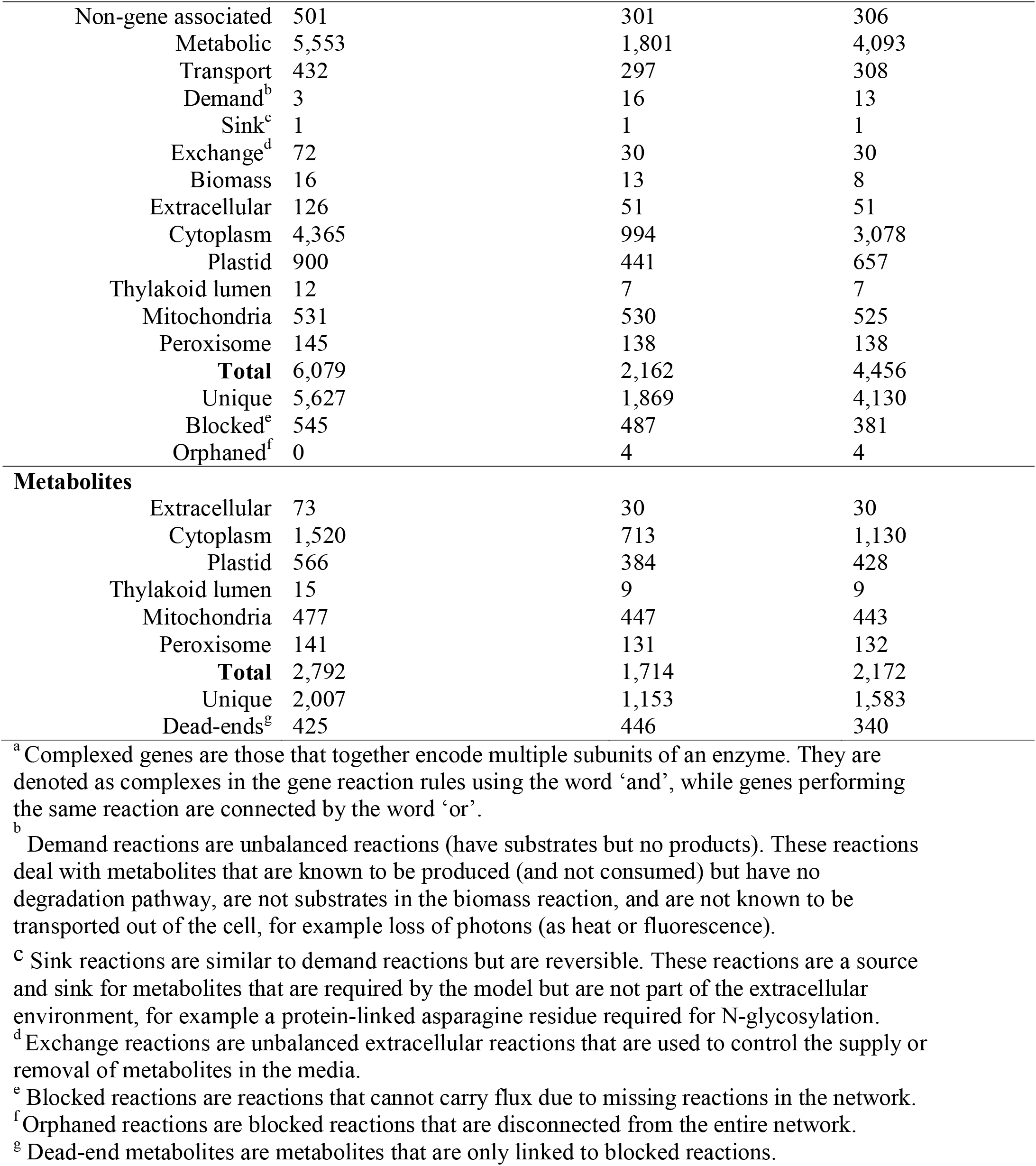
Comparison of attributes of *i*Tps1432 and *P. tricornutum* GEMs

## References

1. Falkowski PG. The Evolution of Modern Eukaryotic Phytoplankton. Science (80-). 2004;305: 354–360. doi:10.1126/science.1095964

2. Sims PA, Mann DG, Medlin LK. Evolution of the diatoms: insights from fossil, biological and molecular data. Phycologia. 2006;45: 361–402. doi:10.2216/05-22.1

3. Allen JF. Photosynthesis of ATP—Electrons, Proton Pumps, Rotors, and Poise. Cell. 2002;110: 273–276. doi:10.1016/S0092-8674(02)00870-X

4. Kramer DM, Evans JR. The Importance of Energy Balance in Improving Photosynthetic Productivity. Plant Physiol. 2011;155: 70–78. doi:10.1104/pp.110.166652

5. Kyle DJ, Ohad I, Arntzen CJ. Membrane protein damage and repair: Selective loss of a quinoneprotein function in chloroplast membranes. Botany. 1984;81: 4070–4074. doi:10.1073/pnas.81.13.4070

6. Asada K. The water–water cycle as alternative photon and electron sinks. Osmond CB, Foyer CH, Bock G, editors. Philos Trans R Soc London Ser B Biol Sci. 2000;355: 1419–1431. doi:10.1098/rstb.2000.0703

7. Munekage Y, Hashimoto M, Miyake C, Tomizawa K-I, Endo T, Tasaka M, et al. Cyclic electron flow around photosystem I is essential for photosynthesis. Nature. 2004;429: 579–582. doi:10.1038/nature02598

8. Allen J. Oxygen reduction and optimum production of ATP in photosynthesis. Nature. 1975;256: 599–600. doi:10.1038/256599a0

9. Bailleul B, Berne N, Murik O, Petroutsos D, Prihoda J, Tanaka A, et al. Energetic coupling between plastids and mitochondria drives CO2 assimilation in diatoms. Nature. 2015;524: 366–369. doi:10.1038/nature14599

10. Dyhrman ST, Benitez-Nelson CR, Orchard ED, Haley ST, Pellechia PJ. A microbial source of phosphonates in oligotrophic marine systems. Nat Geosci. 2009;2: 696–699. doi:10.1038/ngeo639

11. Varma A, Palsson BO. Stoichiometric flux balance models quantitatively predict growth and metabolic by-product secretion in wild-type Escherichia coli W3110. Appl Environ Microbiol. 1994;60: 3724–3731. doi:10.1128/AEM.60.10.3724-3731.1994

12. Klausmeier CA, Litchman E, Levin SA. Phytoplankton growth and stoichiometry under multiple nutrient limitation. Limnol Oceanogr. 2004;49: 1463–1470. doi:10.4319/lo.2004.49.4_part_2.1463

13. Price NM. The elemental stoichiometry and composition of an iron-limited diatom. Limnol Oceanogr. 2005;50: 1159–1171. doi:10.4319/lo.2005.50.4.1159

14. Verschoor A, Van Dijk M, Huisman J, Van Donk E. Elevated CO2 concentrations affect the elemental stoichiometry and species composition of an experimental phytoplankton community. Freshw Biol. 2013;58: 597–611. doi:10.1111/j.1365-2427.2012.02833.x

15. Finkel Z V., Quigg A, Raven JA, Reinfelder JR, Schofield OE, Falkowski PG. Irradiance and the elemental stoichiometry of marine phytoplankton. Limnol Oceanogr. 2006;51: 2690–2701. doi:10.4319/lo.2006.51.6.2690

16. Danger M, Oumarou C, Benest D, Lacroix G. Bacteria can control stoichiometry and nutrient limitation of phytoplankton. Funct Ecol. 2007;21: 202–210. doi:10.1111/j.1365-2435.2006.01222.x

17. Zlotnik I, Dubinsky Z. The effect of light and temperature on DOC excretion by phytoplankton. Limnol Oceanogr. 1989;34: 831–839. doi:10.4319/lo.1989.34.5.0831

18. Lomas MW, Glibert PM. Temperature regulation of nitrate uptake: A novel hypothesis about nitrate uptake and reduction in cool-water diatoms. Limnol Oceanogr. 1999;44: 556–572. doi:10.4319/lo.1999.44.3.0556

19. Myklestad SM. Release of extracellular products by phytoplankton with special emphasis on polysaccharides. Sci Total Environ. 1995;165: 155–164. doi:10.1016/0048-9697(95)04549-G

20. Granum E, Kirkvold S, Myklestad S. Cellular and extracellular production of carbohydrates and amino acids by the marine diatom Skeletonema costatum: diel variations and effects of N depletion. Mar Ecol Prog Ser. 2002;242: 83–94. doi:10.3354/meps242083

21. Durham BP, Boysen AK, Carlson LT, Groussman RD, Heal KR, Cain KR, et al. Sulfonate-based networks between eukaryotic phytoplankton and heterotrophic bacteria in the surface ocean. Nat Microbiol. 2019;4: 1706–1715. doi:10.1038/s41564-019-0507-5

22. Orth JD, Thiele I, Palsson BØ. What is flux balance analysis? Nat Biotechnol. 2010;28: 245–248. doi:10.1038/nbt.1614

23. Fisher NL, Halsey KH. Mechanisms that increase the growth efficiency of diatoms in low light. Photosynth Res. 2016;129: 183–197. doi:10.1007/s11120-016-0282-6

24. Liefer JD, Garg A, Campbell DA, Irwin AJ, Finkel Z V. Nitrogen starvation induces distinct photosynthetic responses and recovery dynamics in diatoms and prasinophytes. Ianora A, editor. PLoS One. 2018;13: e0195705. doi:10.1371/journal.pone.0195705

25. Liefer JD, Garg A, Fyfe MH, Irwin AJ, Benner I, Brown CM, et al. The Macromolecular Basis of Phytoplankton C:N:P Under Nitrogen Starvation. Front Microbiol. 2019;10: 1–16. doi:10.3389/fmicb.2019.00763

26. Levering J, Broddrick J, Dupont CL, Peers G, Beeri K, Mayers J, et al. Genome-Scale Model Reveals Metabolic Basis of Biomass Partitioning in a Model Diatom. Ianora A, editor. PLoS One. 2016;11: e0155038. doi:10.1371/journal.pone.0155038

27. Gruber A, Rocap G, Kroth PG, Armbrust EV, Mock T. Plastid proteome prediction for diatoms and other algae with secondary plastids of the red lineage. Plant J. 2015;81: 519–528. doi:10.1111/tpj.12734

28. Caspi R, Altman T, Billington R, Dreher K, Foerster H, Fulcher CA, et al. The MetaCyc database of metabolic pathways and enzymes and the BioCyc collection of Pathway/Genome Databases. Nucleic Acids Res. 2014;42: D459–D471. doi:10.1093/nar/gkt1103

29. Parker K. Metabolic Network Construction Based on the Genome of the Marine Diatom Thalassiosira Pseudonana and the Analysis of Genome-wide Transcriptome Data to Investigate Triacylglyceride Accumulation. San Jose State University. 2013.

30. Li L. OrthoMCL: Identification of Ortholog Groups for Eukaryotic Genomes. Genome Res. 2003;13: 2178–2189. doi:10.1101/gr.1224503

31. King ZA, Lu J, Dräger A, Miller P, Federowicz S, Lerman JA, et al. BiGG Models: A platform for integrating, standardizing and sharing genome-scale models. Nucleic Acids Res. 2016;44: D515–D522. doi:10.1093/nar/gkv1049

32. Thiele I, Palsson BØ. A protocol for generating a high-quality genome-scale metabolic reconstruction. Nat Protoc. 2010;5: 93–121. doi:10.1038/nprot.2009.203

33. Jones P, Binns D, Chang H-Y, Fraser M, Li W, McAnulla C, et al. InterProScan 5: genome-scale protein function classification. Bioinformatics. 2014;30: 1236–1240. doi:10.1093/bioinformatics/btu031

34. Kanehisa M, Furumichi M, Tanabe M, Sato Y, Morishima K. KEGG: new perspectives on genomes, pathways, diseases and drugs. Nucleic Acids Res. 2017;45: D353–D361. doi:10.1093/nar/gkw1092

35. Elbourne LDH, Tetu SG, Hassan KA, Paulsen IT. TransportDB 2.0: a database for exploring membrane transporters in sequenced genomes from all domains of life. Nucleic Acids Res. 2017;45: D320–D324. doi:10.1093/nar/gkw1068

36. Ebrahim A, Lerman JA, Palsson BO, Hyduke DR. COBRApy: COnstraints-Based Reconstruction and Analysis for Python. BMC Syst Biol. 2013;7: 74. doi:10.1186/1752-0509-7-74

37. Fritzemeier CJ, Hartleb D, Szappanos B, Papp B, Lercher MJ. Erroneous energy-generating cycles in published genome scale metabolic networks: Identification and removal. Maranas CD, editor. PLOS Comput Biol. 2017;13: e1005494. doi:10.1371/journal.pcbi.1005494

38. Hartleb D, Jarre F, Lercher MJ. Improved Metabolic Models for E. coli and Mycoplasma genitalium from GlobalFit, an Algorithm That Simultaneously Matches Growth and Non-Growth Data Sets. Patil KR, editor. PLOS Comput Biol. 2016;12: e1005036. doi:10.1371/journal.pcbi.1005036

39. Broddrick JT, Du N, Smith SR, Tsuji Y, Jallet D, Ware MA, et al. Cross◻compartment metabolic coupling enables flexible photoprotective mechanisms in the diatom Phaeodactylum tricornutum. New Phytol. 2019;222: 1364–1379. doi:10.1111/nph.15685

40. Emanuelsson O, Brunak S, von Heijne G, Nielsen H. Locating proteins in the cell using TargetP, SignalP and related tools. Nat Protoc. 2007;2: 953–971. doi:10.1038/nprot.2007.131

41. Petersen TN, Brunak S, von Heijne G, Nielsen H. SignalP 4.0: discriminating signal peptides from transmembrane regions. Nat Methods. 2011;8: 785–786. doi:10.1038/nmeth.1701

42. Gschloessl B, Guermeur Y, Cock JM. HECTAR: A method to predict subcellular targeting in heterokonts. BMC Bioinformatics. 2008;9: 393. doi:10.1186/1471-2105-9-393

43. Claros MG, Vincens P. Computational Method to Predict Mitochondrially Imported Proteins and their Targeting Sequences. Eur J Biochem. 1996;241: 779–786. doi:10.1111/j.1432-1033.1996.00779.x

44. Cokol M, Nair R, Rost B. Finding nuclear localization signals. EMBO Rep. 2000;1: 411–415. doi:10.1093/embo-reports/kvd092

45. Gattiker A, Gasteiger E, Bairoch A. ScanProsite: a reference implementation of a PROSITE scanning tool. Appl Bioinformatics. 2002;1: 107–8. doi:10.1007/978-1-59745-398-1_25

46. Gonzalez NH, Felsner G, Schramm FD, Klingl A, Maier U-G, Bolte K. A single peroxisomal targeting signal mediates matrix protein import in diatoms. Waller RF, editor. PLoS One. 2011;6: e25316. doi:10.1371/journal.pone.0025316

47. Kroth PG, Chiovitti A, Gruber A, Martin-Jezequel V, Mock T, Parker MS, et al. A model for carbohydrate metabolism in the diatom Phaeodactylum tricornutum deduced from comparative whole genome analysis. Kroymann J, editor. PLoS One. 2008;3: e1426. doi:10.1371/journal.pone.0001426

48. Broddrick JT, Rubin BE, Welkie DG, Du N, Mih N, Diamond S, et al. Unique attributes of cyanobacterial metabolism revealed by improved genome-scale metabolic modeling and essential gene analysis. Proc Natl Acad Sci. 2016;113: E8344–E8353. doi:10.1073/pnas.1613446113

49. McCarthy A, Rogers SP, Duffy SJ, Campbell DA. Elevated carbon dioxide differentially alters the photophysiology of Thalassiosira pseudonana (Bacillariophyceae) and Emiliana huxleyi (Haptophyta). J Phycol. 2012;48: 635–646. doi:10.1111/j.1529-8817.2012.01171.x

50. Shang F. The response of fatty acids and pigments to variations in temperature and irradiance in the Marine Diatom Thalassiosira pseudonana. 2011.

51. Devred E, Turpie K, Moses W, Klemas V, Moisan T, Babin M, et al. Future Retrievals of Water Column Bio-Optical Properties using the Hyperspectral Infrared Imager (HyspIRI). Remote Sens. 2013;5: 6812–6837. doi:10.3390/rs5126812

52. Baird ME, Mongin M, Rizwi F, Bay LK, Cantin NE, Soja-Woźniak M, et al. A mechanistic model of coral bleaching due to temperature-mediated light-driven reactive oxygen build-up in zooxanthellae. Ecol Modell. 2018;386: 20–37. doi:10.1016/j.ecolmodel.2018.07.013

53. Finkel ZV. Light absorption and size scaling of light-limited metabolism in marine diatoms. Limnol Oceanogr. 2001;46: 86–94. doi:10.4319/lo.2001.46.1.0086

54. Stramski D, Sciandra A, Claustre H. Effects of temperature, nitrogen, and light limitation on the optical properties of the marine diatom Thalassiosira pseudonana. Limnol Oceanogr. 2002;47: 392–403. doi:10.4319/lo.2002.47.2.0392

55. Sobrino C, Ward ML, Neale PJ. Acclimation to elevated carbon dioxide and ultraviolet radiation in the diatom Thalassiosira pseudonana◻: Effects on growth, photosynthesis, and spectral sensitivity of photoinhibition. Limnol Oceanogr. 2008;53: 494–505. doi:10.4319/lo.2008.53.2.0494

56. Li G, Campbell DA. Rising CO2 Interacts with Growth Light and Growth Rate to Alter Photosystem II Photoinactivation of the Coastal Diatom Thalassiosira pseudonana. Subramanyam R, editor. PLoS One. 2013;8: e55562. doi:10.1371/journal.pone.0055562

57. Campbell DA, Hossain Z, Cockshutt AM, Zhaxybayeva O, Wu H, Li G. Photosystem II protein clearance and FtsH function in the diatom Thalassiosira pseudonana. Photosynth Res. 2013;115: 43–54. doi:10.1007/s11120-013-9809-2

58. Trimborn S, Wolf-Gladrow D, Richter K-U, Rost B. The effect of pCO2 on carbon acquisition and intracellular assimilation in four marine diatoms. J Exp Mar Bio Ecol. 2009;376: 26–36. doi:10.1016/j.jembe.2009.05.017

59. Platt T, Gallegos C, Harrison W. Photoinhibition of photosynthesis in natural assemblages. J Mar Res. 1980;38.

60. Vedalankar P, Tripathy BC. Evolution of light-independent protochlorophyllide oxidoreductase. Protoplasma. 2019;256: 293–312. doi:10.1007/s00709-018-1317-y

61. Jallet D, Caballero MA, Gallina AA, Youngblood M, Peers G. Photosynthetic physiology and biomass partitioning in the model diatom Phaeodactylum tricornutum grown in a sinusoidal light regime. Algal Res. 2016;18: 51–60. doi:10.1016/j.algal.2016.05.014

62. J. Geider R, Osborne BA, Raven JA. Light dependence of growth and photosynthesis in Phaeodactylum tricornutum (Bacillariophyceae). J Phycol. 2004;21: 609–619. doi:10.1111/j.0022-3646.1985.00609.x

63. Brown MR, Dunstan GA, Norwood SJ, Miller KA. Effects of harvest stage and light on the biochemical composition of the diatom Thalassiosira pseudonana. J Phycol. 1996;32: 64–73. doi:10.1111/j.0022-3646.1996.00064.x

64. Hockin NL, Mock T, Mulholland F, Kopriva S, Malin G. The Response of Diatom Central Carbon Metabolism to Nitrogen Starvation Is Different from That of Green Algae and Higher Plants. Plant Physiol. 2012;158: 299–312. doi:10.1104/pp.111.184333

65. Hunter JE. Phytoplankton Lipidomics◻: Lipid Dynamics in Response to Microalgal Stressors. University of Southampton. 2015.

66. Yu ET, Zendejas FJ, Lane PD, Gaucher S, Simmons BA, Lane TW. Triacylglycerol accumulation and profiling in the model diatoms Thalassiosira pseudonana and Phaeodactylum tricornutum (Baccilariophyceae) during starvation. J Appl Phycol. 2009;21: 669–681. doi:10.1007/s10811-008-9400-y

67. Chiu H, Levy R, Borenstein E. Emergent Biosynthetic Capacity in Simple Microbial Communities. Ouzounis CA, editor. PLoS Comput Biol. 2014;10: e1003695. doi:10.1371/journal.pcbi.1003695

68. Noecker C, Chiu H-C, McNally CP, Borenstein E. Defining and Evaluating Microbial Contributions to Metabolite Variation in Microbiome-Metabolome Association Studies. mSystems. 2019;4: 1–28. doi:10.1128/msystems.00579-19

69. Cermeño P, Marañón E, Romero OE. Response of marine diatom communities to Late Quaternary abrupt climate changes. J Plankton Res. 2013;35: 12–21. doi:10.1093/plankt/fbs073

70. Yin K, Harrison PJ, Dortch Q. Lack of ammonium inhibition of nitrate uptake for a diatom grown under low light conditions. J Exp Mar Bio Ecol. 1998;228: 151–165. doi:10.1016/S0022-0981(98)00025-2

71. Perry MJ. Phosphate utilization by an oceanic diatom in phosphorus-limited chemostat culture and in the oligotrophic waters of the central North Pacific1. Limnol Oceanogr. 1976;21: 88–107. doi:10.4319/lo.1976.21.1.0088

72. Dahlgren B. ChemPy: A package useful for chemistry written in Python. J Open Source Softw. 2018;3: 565. doi:10.21105/joss.00565

73. Ahmad A, Tiwari A, Srivastava S. A Genome-Scale Metabolic Model of Thalassiosira pseudonana CCMP 1335 for a Systems-Level Understanding of Its Metabolism and Biotechnological Potential. Microorganisms. 2020;8: 1396. doi:10.3390/microorganisms8091396

74. Schober AF, Rio Bartulos C, Bischoff A, Lepetit B, Gruber A, Kroth PG. Organelle Studies and Proteome Analyses of Mitochondria and Plastids Fractions from the Diatom Thalassiosira pseudonana. Plant Cell Physiol. 2019;60: 1811–1828. doi:10.1093/pcp/pcz097

75. Helliwell KE, Wheeler GL, Leptos KC, Goldstein RE, Smith AG. Insights into the Evolution of Vitamin B12 Auxotrophy from Sequenced Algal Genomes. Mol Biol Evol. 2011;28: 2921–2933. doi:10.1093/molbev/msr124

76. Heal KR, Qin W, Ribalet F, Bertagnolli AD, Coyote-Maestas W, Hmelo LR, et al. Two distinct pools of B12 analogs reveal community interdependencies in the ocean. Proc Natl Acad Sci. 2017;114: 364–369. doi:10.1073/pnas.1608462114

77. Durkin CA, Mock T, Armbrust EV. Chitin in Diatoms and Its Association with the Cell Wall. Eukaryot Cell. 2009;8: 1038–1050. doi:10.1128/EC.00079-09

78. Montsant A, Jabbari K, Maheswari U, Bowler C. Comparative Genomics of the Pennate Diatom Phaeodactylum tricornutum. Plant Physiol. 2005;137: 500–513. doi:10.1104/pp.104.052829

79. Samukawa M, Shen C, Hopkinson BM, Matsuda Y. Localization of putative carbonic anhydrases in the marine diatom, Thalassiosira pseudonana. Photosynth Res. 2014;121: 235–249. doi:10.1007/s11120-014-9967-x

80. Durham BP, Sharma S, Luo H, Smith CB, Amin SA, Bender SJ, et al. Cryptic carbon and sulfur cycling between surface ocean plankton. Proc Natl Acad Sci. 2015;112: 453–457. doi:10.1073/pnas.1413137112

81. Sarmento H, Gasol JM. Use of phytoplankton-derived dissolved organic carbon by different types of bacterioplankton. Environ Microbiol. 2012;14: 2348–2360. doi:10.1111/j.1462-2920.2012.02787.x

82. Chang RL, Ghamsari L, Manichaikul A, Hom EFY, Balaji S, Fu W, et al. Metabolic network reconstruction of Chlamydomonas offers insight into light◻driven algal metabolism. Mol Syst Biol. 2011;7: 518. doi:10.1038/msb.2011.52

83. La Vars SM, Johnston MR, Hayles J, Gascooke JR, Brown MH, Leterme SC, et al. 29Si{1H} CP-MAS NMR comparison and ATR-FTIR spectroscopic analysis of the diatoms Chaetoceros muelleri and Thalassiosira pseudonana grown at different salinities. Anal Bioanal Chem. 2013;405: 3359–3365. doi:10.1007/s00216-013-6746-z

84. Claquin P, Kromkamp JC, Martin-Jezequel V. Relationship between photosynthetic metabolism and cell cycle in a synchronized culture of the marine alga Cylindrotheca fusiformis (Bacillariophyceae). Eur J Phycol. 2004;39: 33–41. doi:10.1080/0967026032000157165

85. Thamatrakoln K, Hildebrand M. Silicon Uptake in Diatoms Revisited: A Model for Saturable and Nonsaturable Uptake Kinetics and the Role of Silicon Transporters. Plant Physiol. 2008;146: 1397–1407. doi:10.1104/pp.107.107094

86. Kliphuis AMJ, Klok AJ, Martens DE, Lamers PP, Janssen M, Wijffels RH. Metabolic modeling of Chlamydomonas reinhardtii: energy requirements for photoautotrophic growth and maintenance. J Appl Phycol. 2012;24: 253–266. doi:10.1007/s10811-011-9674-3

87. Chalker BE, Dunlap WC, Oliver JK. Bathymetric adaptations of reef-building corals at davies reef, great barrier reef, Australia II. Light saturation curves for photosynthesis and respiration. J Exp Mar Bio Ecol. 1983;73: 37–56. doi:10.1016/0022-0981(83)90004-7

88. Halsey KH, O’Malley RT, Graff JR, Milligan AJ, Behrenfeld MJ. A common partitioning strategy for photosynthetic products in evolutionarily distinct phytoplankton species. New Phytol. 2013;198: 1030–1038. doi:10.1111/nph.12209

89. Jordan DB, Ogren WL. Species variation in the specificity of ribulose biphosphate carboxylase/oxygenase. Nature. 1981;291: 513–515. doi:10.1038/291513a0

90. Young JN, Heureux AMC, Sharwood RE, Rickaby REM, Morel FMM, Whitney SM. Large variation in the Rubisco kinetics of diatoms reveals diversity among their carbon-concentrating mechanisms. J Exp Bot. 2016;67: 3445–3456. doi:10.1093/jxb/erw163

91. Young JN, Hopkinson BM. The potential for co-evolution of CO2-concentrating mechanisms and Rubisco in diatoms. J Exp Bot. 2017;68: 3751–3762. doi:10.1093/jxb/erx130

92. Heureux AMC, Young JN, Whitney SM, Eason-Hubbard MR, Lee RBY, Sharwood RE, et al. The role of Rubisco kinetics and pyrenoid morphology in shaping the CCM of haptophyte microalgae. J Exp Bot. 2017;68: 3959–3969. doi:10.1093/jxb/erx179

93. Shuler M, Kargi F. Stoichiometry of Microbial growth and Product Formation. Bioprocess Eng. 2002; 209–16.

94. Peltier G, Cournac L. Chlororespiration. Annu Rev Plant Biol. 2002;53: 523–550. doi:10.1146/annurev.arplant.53.100301.135242

95. Grouneva I, Muth-Pawlak D, Battchikova N, Aro E-M. Changes in Relative Thylakoid Protein Abundance Induced by Fluctuating Light in the Diatom Thalassiosira pseudonana. J Proteome Res. 2016;15: 1649–1658. doi:10.1021/acs.jproteome.6b00124

96. Cruz S, Goss R, Wilhelm C, Leegood R, Horton P, Jakob T. Impact of chlororespiration on non-photochemical quenching of chlorophyll fluorescence and on the regulation of the diadinoxanthin cycle in the diatom Thalassiosira pseudonana. J Exp Bot. 2011;62: 509–519. doi:10.1093/jxb/erq284

97. Jakob T, Goss R, Wilhelm C. Activation of diadinoxanthin de-epoxidase due to a chlororespiratory proton gradient in the dark in the diatom Phaeodactylum tricornutum. Plant Biol. 1999;1: 76–82. doi:10.1111/j.1438-8677.1999.tb00711.x

98. Davis A, Abbriano R, Smith SR, Hildebrand M. Clarification of Photorespiratory Processes and the Role of Malic Enzyme in Diatoms. Protist. 2017;168: 134–153. doi:10.1016/j.protis.2016.10.005

99. Schnitzler Parker M, Armbrust EV, Piovia-Scott J, Keil RG. Induction of photorespiration by light in the centric diatom Thalassiosira weissflogii (Bacillariophyceae): Molecular characterization and physiological consequences. J Phycol. 2004;40: 557–567. doi:10.1111/j.1529-8817.2004.03184.x

100. Parker MS, Armbrust EV. Synergistic effects of light, temperature, and nitrogen source on transcription of genes for carbon and nitrogen metabolism in the centric diatom Thalassiosira pseudonana (Bacillariophyceae). J Phycol. 2005;41: 1142–1153. doi:10.1111/j.1529-8817.2005.00139.x

101. Turpin DH, Elrifi IR, Birch DG, Weger HG, Holmes JJ. Interactions between photosynthesis, respiration, and nitrogen assimilation in microalgae. Can J Bot. 1988;66: 2083–2097. doi:10.1139/b88-286

102. Bucciarelli E, Sunda WG. Influence of CO2, nitrate, phosphate, and silicate limitation on intracellular dimethylsulfoniopropionate in batch cultures of the coastal diatom Thalassiosira pseudonana. Limnol Oceanogr. 2003;48: 2256–2265. doi:10.4319/lo.2003.48.6.2256

103. Lomas MW. Ammonium release by nitrogen sufficient diatoms in response to rapid increases in irradiance. J Plankton Res. 2000;22: 2351–2366. doi:10.1093/plankt/22.12.2351

104. Rosenwasser S, Graff van Creveld S, Schatz D, Malitsky S, Tzfadia O, Aharoni A, et al. Mapping the diatom redox-sensitive proteome provides insight into response to nitrogen stress in the marine environment. Proc Natl Acad Sci. 2014;111: 2740–2745. doi:10.1073/pnas.1319773111

105. Emerson S, Hedges J. Oceanography background: dissolved chemicls, circulation and biology in the sea. Chemical Oceanography and the Marine Carbon Cycle. Cambridge: Cambridge University Press; 2008. pp. 3–32.

106. Downing J. Marine nitrogen: Phosphorus stoichiometry and the global N:P cycle. Biogeochemistry. 1997;37: 237–252. doi:10.1023%2FA%3A1005712322036

107. Karl D, Letelier R. Nitrogen fixation-enhanced carbon sequestration in low nitrate, low chlorophyll seascapes. Mar Ecol Prog Ser. 2008;364: 257–268. doi:10.3354/meps07547

108. Moore CM, Mills MM, Arrigo KR, Berman-Frank I, Bopp L, Boyd PW, et al. Processes and patterns of oceanic nutrient limitation. Nat Geosci. 2013;6: 701–710. doi:10.1038/ngeo1765

109. Heyer H, Krumbein WE. Excretion of fermentation products in dark and anaerobically incubated cyanobacteria. Arch Microbiol. 1991;155: 284–287. doi:10.1007/BF00252213

110. Husic DW, Tolbert NE. Anaerobic formation of D-lactate and partial purification and characterization of a pyruvate reductase from Chlamydomonas reinhardtii. Plant Physiol. 1985;78: 277–284. doi:10.1104/pp.78.2.277

111. Husic DW, Tolbert NE. Inhibition of glycolate and D-lactate metabolism in a Chlamydomonas reinhardtii mutant deficient in mitochondrial respiration. Proc Natl Acad Sci. 1987;84: 1555–1559. doi:10.1073/pnas.84.6.1555

112. Murik O, Tirichine L, Prihoda J, Thomas Y, Araújo WL, Allen AE, et al. Downregulation of mitochondrial alternative oxidase affects chloroplast function, redox status and stress response in a marine diatom. New Phytol. 2019;221: 1303–1316. doi:10.1111/nph.15479

113. Basan M, Hui S, Okano H, Zhang Z, Shen Y, Williamson JR, et al. Overflow metabolism in Escherichia coli results from efficient proteome allocation. Nature. 2015;528: 99–104. doi:10.1038/nature15765

114. Admiraal W, Peletier H, Laane RWPM. Nitrogen metabolism of marine planktonic diatoms; excretion, assimilation and cellular pools of free amino acids in seven species with different cell size. J Exp Mar Bio Ecol. 1986;98: 241–263. doi:10.1016/0022-0981(86)90216-9

115. Longnecker K, Kido Soule MC, Kujawinski EB. Dissolved organic matter produced by Thalassiosira pseudonana. Mar Chem. 2015;168: 114–123. doi:10.1016/j.marchem.2014.11.003

116. Lau WWY, Armbrust E V. Detection of glycolate oxidase gene glcD diversity among cultured and environmental marine bacteria. Environ Microbiol. 2006;8: 1688–1702. doi:10.1111/j.1462-2920.2006.01092.x

117. Lau WWY, Keil RG, Armbrust EV. Succession and Diel Transcriptional Response of the Glycolate-Utilizing Component of the Bacterial Community during a Spring Phytoplankton Bloom. Appl Environ Microbiol. 2007;73: 2440–2450. doi:10.1128/AEM.01965-06

118. Newton RJ, Griffin LE, Bowles KM, Meile C, Gifford S, Givens CE, et al. Genome characteristics of a generalist marine bacterial lineage. ISME J. 2010;4: 784–798. doi:10.1038/ismej.2009.150

119. Howard EC, Sun S, Biers EJ, Moran MA. Abundant and diverse bacteria involved in DMSP degradation in marine surface waters. Environ Microbiol. 2008;10: 2397–2410. doi:10.1111/j.1462-2920.2008.01665.x

120. Teeling H, Fuchs BM, Becher D, Klockow C, Gardebrecht A, Bennke CM, et al. Substrate-Controlled Succession of Marine Bacterioplankton Populations Induced by a Phytoplankton Bloom. Science (80-). 2012;336: 608–611. doi:10.1126/science.1218344

121. Armbrust E V. The genome of the diatom Thalassiosira Pseudonana: Ecology, evolution, and metabolism. Science (80-). 2004;306: 79–86. doi:10.1126/science.1101156

122. Dyhrman ST, Jenkins BD, Rynearson TA, Saito MA, Mercier ML, Alexander H, et al. The Transcriptome and Proteome of the Diatom Thalassiosira pseudonana Reveal a Diverse Phosphorus Stress Response. Santos P, editor. PLoS One. 2012;7: e33768. doi:10.1371/journal.pone.0033768

123. Ashworth J, Coesel S, Lee A, Armbrust E V., Orellana M V., Baliga NS. Genome-wide diel growth state transitions in the diatom Thalassiosira pseudonana. Proc Natl Acad Sci. 2013;110: 7518–7523. doi:10.1073/pnas.1300962110

124. Valenzuela JJ, López García de Lomana A, Lee A, Armbrust E V., Orellana M V., Baliga NS. Ocean acidification conditions increase resilience of marine diatoms. Nat Commun. 2018;9: 2328. doi:10.1038/s41467-018-04742-3

125. Bender SJ, Durkin CA, Berthiaume CT, Morales RL, Armbrust EV. Transcriptional responses of three model diatoms to nitrate limitation of growth. Front Mar Sci. 2014;1: 1–15. doi:10.3389/fmars.2014.00003

126. Bertrand EM, McCrow JP, Moustafa A, Zheng H, McQuaid JB, Delmont TO, et al. Phytoplankton–bacterial interactions mediate micronutrient colimitation at the coastal Antarctic sea ice edge. Proc Natl Acad Sci. 2015;112: 9938–9943. doi:10.1073/pnas.1501615112

127. Sañudo-Wilhelmy SA, Gómez-Consarnau L, Suffridge C, Webb EA. The Role of B Vitamins in Marine Biogeochemistry. Ann Rev Mar Sci. 2014;6: 339–367. doi:10.1146/annurev-marine-120710-100912

## References

Armbrust E V. The Genome of the Diatom Thalassiosira Pseudonana: Ecology, Evolution, and Metabolism. Science (80-). 2004;306: 79–86. doi:10.1126/science.1101156

Bowler C, Allen AE, Badger JH, Grimwood J, Jabbari K, Kuo A, et al. The Phaeodactylum genome reveals the evolutionary history of diatom genomes. Nature. 2008;456: 239–244. doi:10.1038/nature07410

Broddrick JT, Du N, Smith SR, Tsuji Y, Jallet D, Ware MA, et al. Cross◻compartment metabolic coupling enables flexible photoprotective mechanisms in the diatom Phaeodactylum tricornutum. New Phytol. 2019;222: 1364–1379. doi:10.1111/nph.15685

Gruber A, Rocap G, Kroth PG, Armbrust EV, Mock T. Plastid proteome prediction for diatoms and other algae with secondary plastids of the red lineage. Plant J. 2015;81: 519–528. doi:10.1111/tpj.12734

Levering J, Broddrick J, Dupont CL, Peers G, Beeri K, Mayers J, et al. Genome-Scale Model Reveals Metabolic Basis of Biomass Partitioning in a Model Diatom. Ianora A, editor. PLoS One. 2016;11: e0155038. doi:10.1371/journal.pone.0155038

Schober AF, Río Bártulos C, Bischoff A, Lepetit B, Gruber A, Kroth PG. Organelle Studies and Proteome Analyses of Mitochondria and Plastids Fractions from the Diatom Thalassiosira pseudonana. Plant Cell Physiol. 2019;60: 1811–1828. doi:10.1093/pcp/pcz097

